# The ortholog of the Human T2D Risk Gene *STEAP2* is Required for Glucose Stimulated Insulin Secretion in Mice

**DOI:** 10.64898/2026.09.26.754731

**Authors:** Avery Hawthorne, Kai Kelly, Mark Tigue, Hilana El-Mekkoussi, Eric Waite, Yelda Erhan, Elisabetta Manduchi, Sepideh Dashti, Andrea Rozo, Jeffrey Roman, Nicolai Doliba, Doris Stoffers, Klaus H. Kaestner

## Abstract

Type 2 diabetes is a complex metabolic disease with a strong genetic component, yet this genetic risk is driven by common variants in up to 600 loci with small effect sizes. Variants in the *STEAP2* locus, encoding ‘Six-transmembrane epithelial antigen of the prostate-2’, are highly associated with altered glycemic traits and type 2 diabetes risk. To elucidate its function in pancreatic β-cells, we utilized Crisp-Cas9 assisted homologous recombination to derive a conditional null allele for *Steap2* (*Steap2^fl/fl^*) in mice and crossed this to the rat insulin promoter (Rip)-Cre transgene. Ablation of *Steap2* induced glucose intolerance, blunted glucose-stimulated Ca^2+^ mobilization and stunted first phase glucose-simulated insulin secretion. In contrast, the incretin effect and second phase insulin secretion were not affected. Islet respirometry and transmission electron microscopy (TEM) identified disrupted cristate morphology and impaired glucose-stimulated oxygen consumption as causally responsible for the observed, impaired triggering pathway of insulin secretion. These findings suggest that at least part of the genetic risk for T2D conferred by *Steap2* variants is likely due to its function in maintaining optimal mitochondrial respiration in pancreatic β-cells.

**Article Highlights:**

- Common variants in the *STEAP2* gene are strongly associated with altered glycemic traits and type 2 diabetes.
- Loss of *Steap2* in β-cells impairs mitochondrial respiration, resulting in attenuated glucose-stimulated Ca^2+^ mobilization and insulin secretion.

## Introduction

Type 2 diabetes (T2D) comprises 90–95% of diabetes diagnoses and is characterized by hyperglycemia caused by the inability of insulin producing β-cells to compensate for insufficient insulin signaling which then attenuates insulin-stimulated glucose disposal and fails to suppress hepatic gluconeogenesis (1). Obesity increases the risk of diabetes by 3- to 8-fold, and 85% of individuals with type 2 diabetes are obese (2–5). However, this association is confounded by the fact that while 40% of obese individuals present with insulin resistance, most do not develop diabetes, and many individuals exhibit muscle insulin resistance in the absence of either obesity or diabetes. This suggests that additional individual risk factors contribute to the progression from insulin resistance to β-cell insufficiency and type 2 diabetes (6–9). While twin studies have estimated the heritability of T2D to be between 31–72% (10; 11), monogenic causes of diabetes—which include (MODY)—implicate at most 60 genes and contribute to only ∼5% of diabetes cases (12).

Genome wide association studies (GWAS) have implicated over 1,200 lead SNPs at approximately 600 loci, each with relatively small effects on T2D disease risk (OR 1.05–1.1), with 200 loci sufficient to explain approximately 25% of T2D heritability (13; 14). Efforts by the NIDDK-funded Advancing Medical Partnerships (AMP) program to integrate pre-existing GWAS studies with new functional datasets have produced the CMD Knowledge Portal (15), which utilizes HuGE scores (16) to quantify the genetic support for involvement of a gene in phenotypic traits. Among the genes deemed to have HuGE scores ranging from “strong” to “compelling” for various glycemic traits, including type 2 diabetes, is the metalloreductase Six-Transmembrane Epithelial Antigen of Prostate 2 (*STEAP2*). At present, it is not known how *STEAP2* affects glycemic control.

Steap2 belongs to the family of STEAP metalloreductases, which reduce Fe³⁺ to Fe²⁺ to facilitate iron transport by the divalent metal transporter 1 (DMT1) from endocytic vesicles into the cytoplasm, where iron is either stored in ferritin (Ft) or transported to the mitochondria for incorporation into Fe–S clusters and heme (17–19). These processes are necessary for the metalation of electron transport chain (ETC) and tricarboxylic acid (TCA) cycle enzymes, and thus mitochondrial function (19). Serum ferritin levels have been implicated as a risk factor for diabetes and elevated iron confers a four-fold increased risk of T2D onset (20; 21). β-cells are uniquely susceptible to iron overload due to their high expression of *DMT1* and *Zip14* which import Fe^3+^ and are required for Zn²⁺ import during insulin granule packaging in conjunction with low expression of the iron exporter ferroportin and ROS-detoxifying enzymes such as superoxide dismutase (SOD) and catalase (22–26). This arrangement likely exists because β-cells engage in a high degree of iron metabolism to fulfill the extensive demand for enzyme metalation required for mitochondrial function during glucose sensing (27). In mice, TFR1-mediated iron uptake is necessary for post-natal β-cell maturation and survival in early development but is dispensable in adulthood (28). It is possible that in later stages of life iron pools are sufficiently maintained through alternative non-transferrin bound iron (NTBI) uptake pathways, as absolute iron deprivation through chelation attenuates glucose-stimulated insulin secretion (GSIS) in adult, murine isolated islets and disrupts the expression of β-cell identity genes (27; 29). Interestingly, β-cell specific deletion of *Dmt1* has little effect on glucose tolerance or *in vitro* GSIS (30), suggesting that NTBI uptake from endosomes is primarily mediated by Zip14 in concert with STEAPs.

To date, it remains unclear whether *Steap2* serves as the primary β-cell metalloreductase and whether its role in iron import is critical for the metalation of enzymes necessary for insulin synthesis and glucose-stimulated insulin secretion. *Steap4* plays a central role in adipocyte progenitor differentiation, and its depletion induces profound insulin resistance (31; 32). To clarify the role of Steap2 in glycemic control, we generated a mouse model of embryonic β-cell-specific deletion of *Steap2* to investigate its function in β-cell insulin secretion dynamics.

## Research Design and Study Methods

### Derivation of Transgenic Mice

Derivation of the *Steap2^fl/fl^* allele was performed by CRISPR/Cas9 directed homologous recombination in zygotes to introduce *loxP* sites upstream of exon 1 and within intron 5 of the *Steap2* gene. The sequences of the guide RNAs (gRNAs) and repair template are listed in Supplementary Table 1.

### Animals and Housing

All animal procedures were approved by the IACUC of the University of Pennsylvania. Both normal chow (NC) and high fat diet (HFD) cohort animals were housed in cages containing no more than 5 mice and were maintained on a 12-hour (6 AM-6 PM) light/dark cycle. The HFD cohort of mice were placed on the AIN-93G “Western Diet” (Test Diet, Ref#5TJN) at 6 weeks of age and housed in thermoneutral cabinets held at 30°C degrees. Experimental mice were generated by crossing C57bl/j *Steap2^fl/fl^* mice with C57bl/j Rat insulin promoter (Rip) Cre recombinase (Cre) mice (33) to obtain *Steap2^fl/fl^;RipCre^+/-^* mice, referred to as “mutant” (mut). Transgenic mice were genotyped by PCR on distal phalanx toe clippings using the primers and cycling conditions listed in Supplementary Table 2. Random fed glucose and weight were assessed once per week between 9-11 AM from 6-18 weeks of age.

### Glucose and Insulin Tolerance Tests

16 hour-fasted mice were either injected or orally gavaged with 20% D-glucose (RPI, Ref#G32045-500.0) resuspended in sterile ddH_2_O interperitoneally and administered 2 g glucose/kg body weight for normal chow mice and 1 g glucose/kg bodyweight for HFD mice. Blood glucose was measured with an AlphaTrak3 glucometer (Zoetis, Ref#ATK-00309R3) after tail vein bleed. Blood for insulin measurements was collected by submandibular cheek bleed at 0, 2, or 5 min. During continuous telemetric monitoring, blood was collected by tail vein bleed at 0, 5, and 15 min. The plasma fraction was centrifuged in Microvette® CB300 EDTA K2E tubes (Sarstedt, Ref# 20.1339.100) at 2,000 RCF for 30 min at 4°C, and total plasma insulin concentration was detected by ELISA (Crystal Chem, Ref#90080). Insulin tolerance tests were performed on random fed mice at 9 AM by injecting insulin (Lantus, Ref#191.49250.3) (1U/kg body weight).

### Continuous Telemetric Monitoring of HFD mice

6-week-old male mice were fed Western Diet and housed at thermoneutrality for 18 weeks. At 24 weeks, an HD-XG (Data Sciences International, St. Paul, MN) continuous blood glucose sensor was surgically implanted in the aorta and the transmitter was placed subcutaneously in the left flank (34). Mice were single housed for one week before surgery and thereafter, and the cages were placed above the RPC-1 receivers (Data Sciences International, St. Paul, MN) in a thermoneutral cabinet held at 30°C degrees on 12-hour light/dark cycle. Data were relayed by the probes every minute and collected for 3 weeks after recovery. 16-hour-fasted IP-GTTs and O-GTTs (1 g D-glucose/kg bodyweight) were performed twice, and tail vein blood was collected at 0, 5, and 15 min. 16-hour fasts followed by 2-hour refeeds were performed twice, and the results presented are the average of these replicates. The experimental timeline is illustrated in Fig 4A.

### Islet Isolation and Culture

Mouse islets were isolated as described (35). Islets were cultured at 37°C and 5% CO_2_ for 1-2 days in RPMI 1640 (Invitrogen, Ref#11875085) containing 10% fetal bovine serum, 1% penicillin-streptomycin (Gibco, Ref#15140122), 11 mM glucose, 2 mM Glutamine, 1 mM Sodium Pyruvate (Invitrogen, Ref#11360070), 10 mM HEPES (Invitrogen, Ref#15630080).

### Histology and Immunofluorescence

Pancreata were isolated from mice and fixed with 10% neutral buffered formalin (Sigma-Aldrich, Ref#HT5011-1CS) for 24 hours, and fixed tissues in 70% EtOH were embedded in paraffin and sectioned at 5 µm thickness by the Penn Molecular Pathology and Imaging Core. Immunofluorescence staining was performed as described (36). Antibodies are listed in Supplementary Table 3. Images were captured via fluorescence microscopy at 40x magnification with a Keyence BZ-X800 instrument. Insulin, glucagon and somatostatin positive areas were quantified in QuPath to calculate total islet area and islet endocrine cell composition (version 0.5.1.).

### RNA-scope in Situ Hybridization

RNA-scope for *Steap2* mRNA was performed on FFPE-fixed, RNase free, 5 μm thick sections followed by combined immunofluorescent staining for insulin and glucagon as described (37). Probes are listed in Supplementary Table 3. All images were captured on a STELLARIS 5 Confocal Microscope at 40x magnification.

### Transmission Electron Microscopy (TEM)

Dissected pancreata were cut into 2x2 mm blocks and fixed with 2.5% glutaraldehyde, 2.0% paraformaldehyde in 0.1M sodium cacodylate buffer (pH = 7.4) overnight at 4℃ for transmission electron microscopic examination and post-fixed in 2.0% osmium tetroxide with 1.5% K_3_Fe(CN)_6_ for 1 hour prior to staining with 2% uranyl acetate. After dehydration through a graded ethanol series, the tissue was infiltrated and embedded in EMbed-812 (Electron Microscopy Sciences, Fort Washington, PA). Thin sections were stained with uranyl acetate and SATO lead and examined with a Talos L120C electron microscope fitted with a Ceta 16M digital camera.

### Quantitative Polymerase Chain Reaction (qPCR)

Genomic DNA was extracted from isolated islets with Qiagen ATL lysis buffer (Ref# 19076) and Proteinase K (New England Biolabs, Ref#P8107S) followed by phenol-chloroform extraction with Phenol:Chloroform:Isoamyl Alcohol (25:24:1, v/v) (Fisher Scientific, Ref#15-593-031), ethanol precipitation and membrane dialysis with a 0.025 µm MCE membrane filter (Milipore, Ref#VSWP02500) to eliminate excess salt content. qPCR was performed on an Applied Biosystems QuantStudio™ 5 instrument using Syber Green PowerUp Plus (Applied Biosystems, RefA25742) master mix. Primers amplifying β-tubulin and glyceraldehyde-3-phosphate dehydrogenase serving as controls. Likewise, relative mitochondrial copy number was assessed with primers amplifying the mitochondrial gene *MT-RNR2* with the nuclear gene β2-microglobulin serving as control as described (38). See details in Supplementary Tables 1 and 2.

### Perifusion of Islets for Measurement of Insulin Secretion Dynamics

A minimum of 100 islets from each mouse (12-18 weeks old) were handpicked and perifused with the Biorep perifusion system (BioRep Diabetes, Ref#PERI5-115/230) at a flow rate of 200 mL/min with Krebs-Ringer bicarbonate buffer and secretagogues were added to the perfusate reservoir at the time points indicated in Fig 5B. Samples were collected at 1-minute intervals, and total insulin was measured by ELISA (Crystal Chem, Ref#90080). Perfused islets were resuspended and lysed in 200 mM glycine, 0.5% NP40, 1x cOmplete™ Protease Inhibitor Cocktail (Roche, Ref# 04693116001) lysis buffer (pH 8.8) and sonicated for 6 min 30s on/off with the Bioruptor Standard (Diagenode, Model UCD-200TM-EX). Islet lysate protein was quantified by Bradford assay (Sigma-Aldrich, Ref#B6916). Lysate total insulin, mature insulin, and proinsulin content were measured as described above.

### Intracellular Ca^2+^ Measurements by Confocal Imaging

Mouse islets were loaded with Fura-2 AM Ca^2+^ fluorophore in Kreb’s buffer for 40 min at 37°C. Islets were transferred to a perifusion chamber placed on a homeothermic platform maintained at 37°C. Islets were then perfused at a flow rate of 1 ml/min with Krebs-Ringer bicarbonate buffer at 37°C with secretagogues indicated in Fig 5A. Intracellular Ca^2+^ was measured every 10s by dual-wavelength fluorescence with excitation at 340 nm and 380 nm and emission at 510 nm using a Zeiss AxioVision system (36).

### Islet Respirometry by Seahorse

Islets were isolated from the normal chow cohort of mice (12-18 weeks old) or from 30-week-old male mice from the HFD cohort and cultured in islet media for 48 h before each experiment. Islets were washed twice in 0.5% FBS, Seahorse XF DMEM media (Agilent Technologies, Ref#103335-100) and seeded into a poly-L-lysine coated (0.1mg/mL, ddH2O) XF96 spheroid plate (Agilent Technologies, Ref#102959-100) containing 175 μL of 0.5% FBS, 1 mM D-glucose Seahorse XF base medium via gravity. Islets were placed in a non-CO_2_ incubator for 1 hour before the experiment. Oxygen consumption was measured using an Agilent Seahorse XFe96 Analyzer. Basal respiration was measured, followed by the stepwise addition of 16.7 mM glucose, 5 µM oligomycin, 2 µM FCCP, and then 5 µM of rotenone and antimycin A. Oxygen consumption measurements were normalized to islet equivalents (IEQs) determined by imaging in a Cytation 5 Cell Imaging Multimode Imager (Biotek/Agilent Technologies, Santa Clara, CA) (39). 65 µL of 4x lysis buffer consisting of 200 mM Tris-HCl, 40 mM EDTA and 8% SDS was added to islet-containing wells, and DNA was isolated by phenol-chloroform extraction and ethanol precipitation for subsequent mtDNA quantification by qPCR.

### Bulk RNA Sequencing and Analysis of Isolated Islets

RNA was extracted from islets from 8-week-old mice using the Trizol™ Reagent (Thermo Fisher Scientific, Ref#15596026). Samples were sequenced by Azenta Biosciences after a rRNA depletion strategy and DNAase treatment. Paired-end sequencing to 150 bp of UMI-tagged cDNA fragments was performed on an Illumina® NovaSeq™ 6000 instrument with a median read pair count of ∼45M per sample. Sequence reads were trimmed and UMI-based de-duplication performed using fastp v.0.23.1 (40). Reads were then mapped to the *Mus musculus* GRCm38 ERCC reference genome using STAR aligner v.2.5.2b (41). Unique gene hit counts were calculated by using featureCounts from the Subread package v.1.5.2 (42).

To compare control (*Steap2^fl/fl^;RipCre^-/-^*, n=7) to mutant mice (*Steap2^fl/fl^;RipCre^+/-^*, n=5), raw counts were input into DESeq2 v1.42.0 (43). Genes with at least 10 counts in at least 3 samples were kept for analyses (21,478 genes) and DESeq2 was run with design = ∼ sex + condition. GSEA (44) was run in pre-ranked mode with ranking based on the DESeq2 statistic, separately on the mouse Hallmark, GO Biological Process, and Reactome collections.

### Data Analysis

Data (other than bulk RNA-Seq) are presented as mean ± SEM and were analyzed using GraphPad Prism 11 software (version 10.5.0) and RStudio (version 2026.01.2+418). Unpaired, two-tailed Student’s t-tests were employed when comparing the means of two groups. One-way analysis of variance (ANOVA) with post hoc Tukey test was employed when more than two groups were compared with one independent variable, while two-way ANOVA was employed when two independent variables were present. Values were considered statistically significant when p < 0.05.

### HPAP scRNA-Seq data

Human Pancreatic Analysis Program (HPAP) scRNA-Seq data pre-processing and cell annotation are described in (45). Pseudobulk counts for each cell type were generated for each donor with at least 50 cells of that type. Counts were then normalized with DESeq2 across all donors and cell types.

### Data and Resource Availability

The RNAseq data are available from NCBI under accession number GEO xxx. All other data generated or analyzed during this study are included in the published article (and its online supplementary files). The *Steap2*^fl^ mice will be made available via the Mutant Mouse Resource and Research Centers (https://www.mmrrc.org).

## Results

### Derivation of a pancreatic β-cell specific Steap2 null allele

We sought to identify genes in which variants are strongly associated with altered glycemic traits and T2D. Human genetic evidence scores (HuGE) implicated altered fasting glucose, random fed glucose, type 2 diabetes and Hba1C as having compelling to strong evidence for association with variants in the *STEAP2* locus **(Fig 1A)**. According to the Human Pancreatic Analysis Program (HPAP)’s (46; 47) single cell RNA-seq dataset, *STEAP2* mRNA is most abundant in β-cells **(Fig S1A)**.

**Figure 1:**
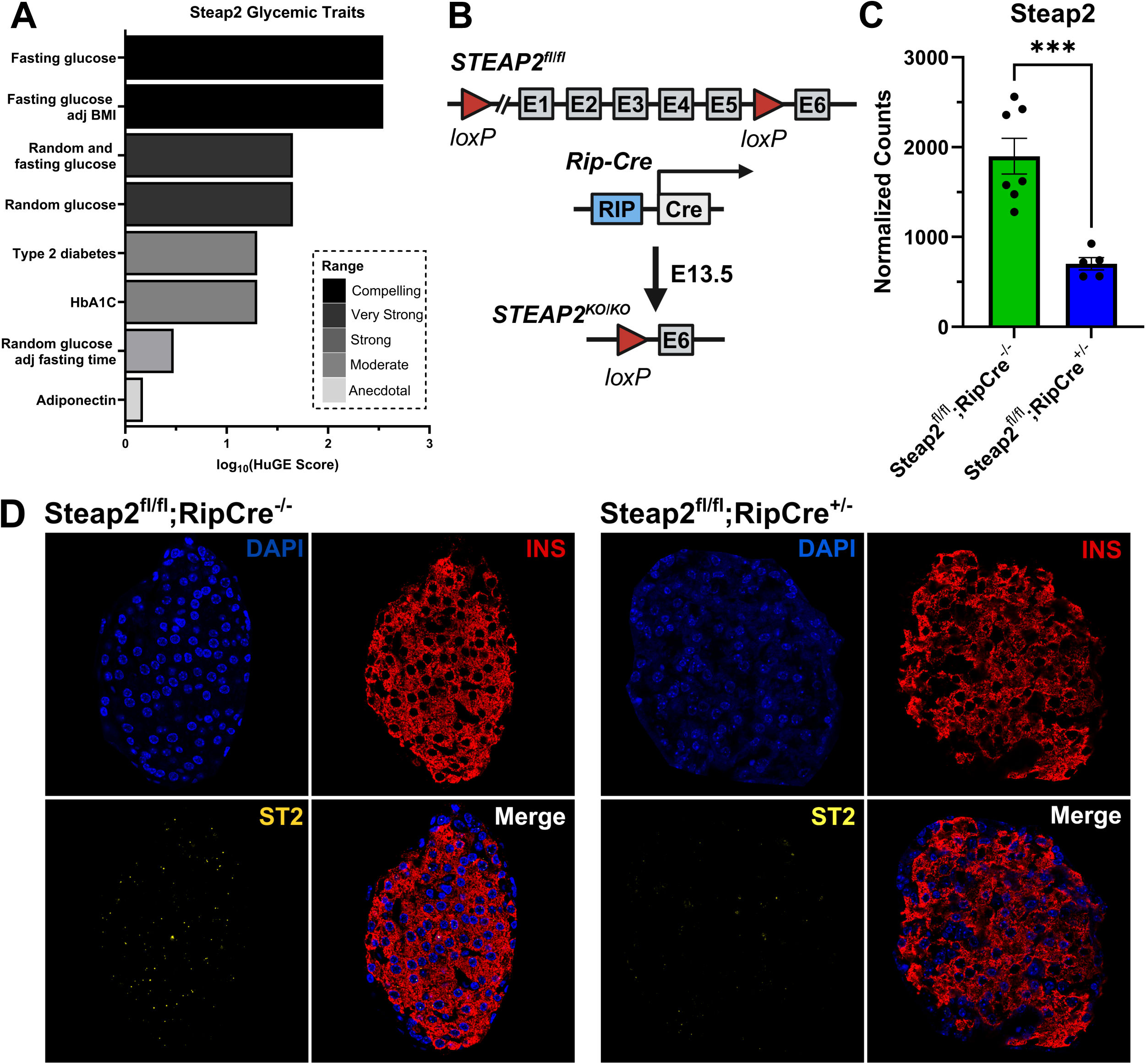
Validation of β-cell specific ablation of *Steap2* in *Steap2^fl/fl^;RipCre^+/-^* mice. **(A)** Summary of glycemic traits associated with variants in *STEAP2*, with their log_10_(HuGE) scores indicated. The rating of genetic support for involvement of *STEAP2* in these traits is denoted by the bar color. **(B)** Design of the *Steap2*^fl/fl^ allele. Experimental (mutant) mice are heterozygous for the β-cell specific, constitutive Rat insulin promoter (Rip)-Cre recombinase, producing *Steap2* deletion in β-cells beginning at E13.5. **(C)** Bulk RNA-Seq in isolated islets from 12–18-week-old mice (Ctrl n=7, Mut n=5). DESeq2 normalized counts confirm ∼64% reduction in *Steap2* mRNA expression (mean ± SEM shown). The DESeq2 p-value = 7.02904^-24^ and the adjusted p-value = 1.62257^-21^. **(D)** RNAScope® fluorescent *in-situ* hybridization (FISH) for *Steap2* mRNA with accompanying insulin and DAPI immunofluorescence staining. Islet morphology is similar between groups, but *Steap2* puncta (yellow) are absent in insulin positive regions in *Steap2^fl/fl^;RipCre^+/-^* mice.

We utilized Crispr-Cas9-assisted homologous recombination in mouse zygotes to introduce *loxP* sites ∼1,000 bp upstream the transcription start site of exon 1 and within intron 5 to derive a conditional null allele for *Steap2* (*Steap2^fl/fl^*) **(Fig 1B)**, ensuring a functional null allele following Cre recombinase (Cre)-dependent recombination (17; 48; 49). Critically, the promoter and ORF for the evolutionarily conserved cilia and flagella associated protein 69 (*Cfap69*), which are upstream of exon 1 and play a critical role in flagella and cilia assembly, are also unaffected. Rat insulin promoter (Rip)-Cre mice (33) were crossed into the *Steap2^fl/fl^* line to produce *Steap2^fl/fl^;RipCre^+/-^*mice (thereafter termed “mutant” and abbreviated to “Mut”). *Steap2^fl/fl^* mice without the RipCre transgene (*Steap2^fl/fl^;RipCre^-/-^*) were used as control, referred to as “control” and abbreviated to “Ctrl”.

qPCR of genomic DNA from isolated islets confirmed that the *Steap2* genomic abundance was reduced by 50% in mutant mice compared to controls **(Fig S1B)**, and *Steap2* mRNA expression was reduced by 64% in the bulk RNA-seq data set **(Fig 1C)**. In addition, we performed RNA fluorescence in situ hybridization (FISH) with RNAscope probes targeting the *Steap2* mRNA in conjunction with immunofluorescence staining for insulin and DAPI to determine the extent of β-cell specific deletion. Control mice exhibited abundant *Steap2* signals within and outside the insulin positive region, while these puncta were largely absent in insulin-positive cells of *Steap2* mutant mice, confirming efficient ablation of *Steap2* (**Fig 1D**).

### Homeostatic glycemic control, weight and islet morphology and composition are not altered by β-cell Steap2 deficiency

Measurements of weight between 6-18 weeks indicated no significant differences between mutant and control mice within both male and female cohorts **(Fig 2A)**. Likewise, random fed glucose during the same time-period was also not significantly different in either sex (**Fig 2B-C**). Immunofluorescence staining for DAPI, insulin, glucagon and somatostatin in pancreas sections did not reveal any gross morphological differences between control and mutant mice. Moreover, pancreas mass normalized to body weight was unchanged (**Fig 2E**). Next, we quantified the average islet area and the percentage of the total islet that was comprised of insulin, glucagon and somatostatin positive regions, but found no significant differences (**Fig 2F**) between groups. We quantified a minimum of 20 islets per mouse in control (n = 5) and mutant (n = 4) mice between 12-18 weeks of age.

**Figure 2:**
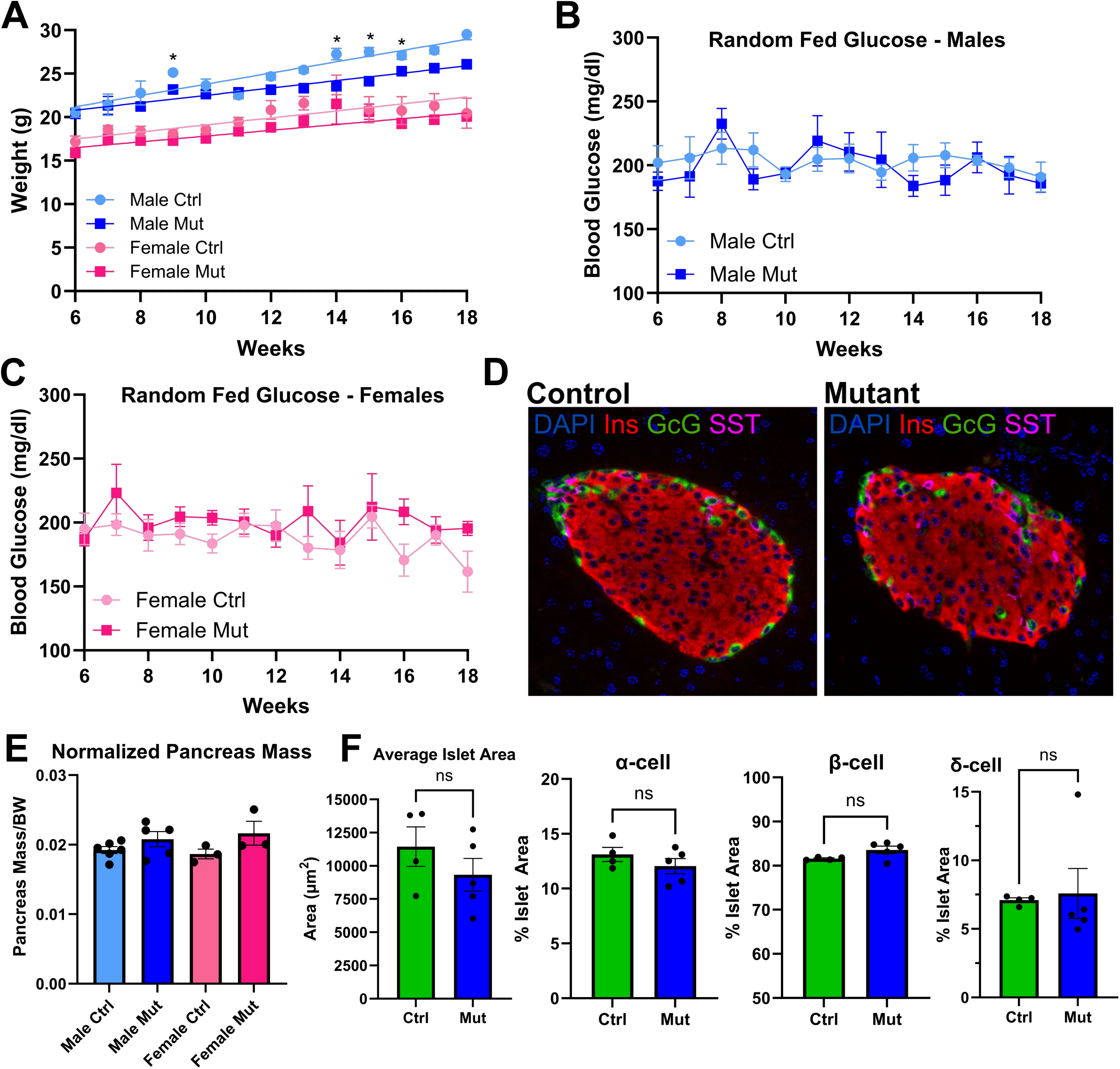
Baseline glycemic control and islet morphology are not different between control and *Steap2* mutant mice. **(A)** Weights of *Steap2* control and mutant mice (n=5) aged 6 to 18 weeks with accompanying slope. No sustained weight differences between sex-matched control and mutant mice were observed. **(B-C)** Random fed glucose of sex-matched, control and mutant mice aged 6-18 weeks (control n = 4, mutant n = 7), showing no differences in blood glucose. **(D)** Immunofluorescence staining of control and mutant mice for insulin (Ins) glucagon (Gcg) and somatostatin (SST). **(E)** Pancreas mass normalized to body weight of sex-matched,12-18-week-old control and mutant mice, showing no differences between sex-matched control and mutant mice. **(F**) Quantification of immunofluorescence staining of average islet area and the percent islet composition of each islet endocrine cell in control (n = 4 mice) and mutant (n = 5 mice) from 5 mm FFPE pancreas sections. A minimum of 20 islets were quantified per mouse, and no differences were observed between sex-matched control and mutant mice. P-values in panels A-E and panel F were separately determined by two-way ANOVA with post-hoc Tukey test and Student’s T-Test, respectively. *, p<0.05, ns = not significant. Mean ± SEM are shown.

### Steap2 mutant mice exhibit glucose intolerance due to attenuated glucose stimulated insulin secretion

Next, we assessed glucose tolerance in mutant mice by 16-hour fasted intraperitoneal (IP-) glucose tolerant test (GTT) in 6-week-old, male (**Fig 3A-B**) and female (**Fig 3D-E**) mice. Following glucose injection, there was a larger glucose excursion in mutant male and mutant female mice compared to their age matched controls, with a larger effect in female mice (**Fig 3A, 3D**). Likewise, total glucose egress quantified by the area under the curve (AUC) was significantly greater in mutant male mice and mutant female mice compared to their age-matched controls, and this difference was greater in female mice (**Fig 3B, 3E**). Insulin Cre lines (such as the MipCre and RipCre) can present with a glucose intolerance phenotype independently of transgene recombination (50–53). We performed an IP-GTT in male wild type (WT;RipCre-) and WT;RipCre^+/-^ mice and found no significant differences in glucose at any timepoints nor any difference in the AUC (**Fig S1C-D**).

**Figure 3:**
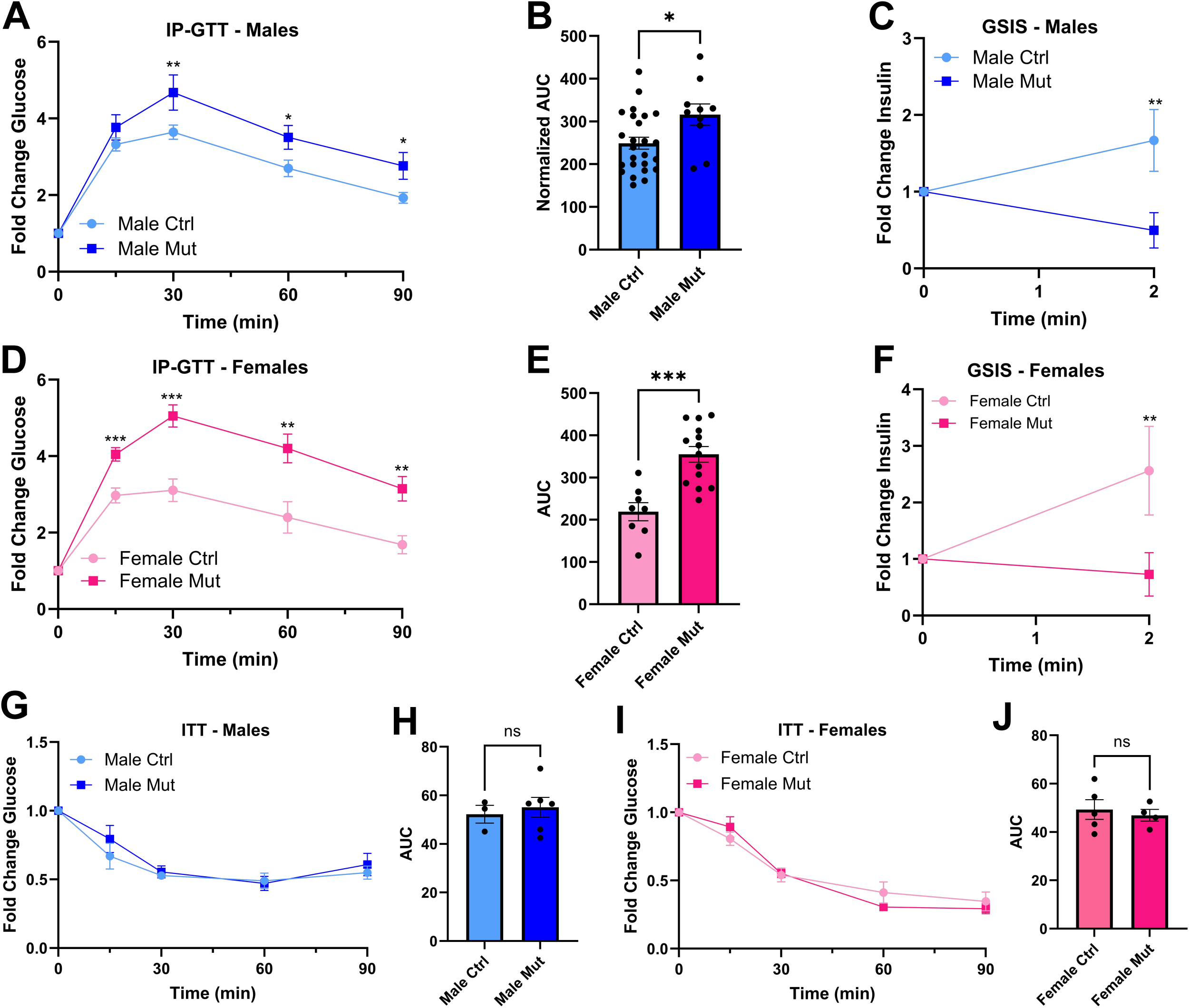
*Steap2* mutant mice exhibit glucose intolerance and attenuated glucose stimulated insulin secretion. **(A)** Intraperitoneal (IP) glucose tolerance tests (GTT) of control (n = 8) and mutant (n = 25) male mice aged 6-7 weeks demonstrated a prolonged and greater glucose excursion in mutant mice. **(B)** Area under the curve (AUC) analysis shows a significant increase in glucose excursion in *Steap2*-deficient male mutant mice. **(C)** In *vivo* GSIS of male control and mutant mice (n = 9) aged 6-7 weeks during an IP-GTT. Glucose-stimulated insulin secretion is significantly decreased in male mutant mice. **(D)** IP-GTT of control (n = 8) and mutant (n = 14) female mice aged 6-7 weeks demonstrated a prolonged and greater glucose excursion in *Steap2*-deficient female mutant mice. **(E)** AUC analysis shows a significant increase in glucose excursion in female mutant mice. **(F)** *In vivo* GSIS of female control and mutant mice (n = 9) aged 6-7 weeks during an IPGTT. Glucose-stimulated insulin secretion is significantly decreased in female mutant mice. **(G)** Insulin tolerance tests (ITT) in 6-to 7-week-old male control (n = 3) and mutant (n = 6) mice demonstrating indistinguishable insulin sensitivity. **(H)** AUC analysis of the ITT shows no significant differences in insulin action. **(I)** Insulin tolerance tests (ITT) in 6- to 7-week-old female control (n = 5) and mutant (n = 4) mice demonstrating indistinguishable insulin sensitivity. **(J)** AUC analysis of the ITT demonstrating no significant differences in insulin action. P-values for IP-GTT, ITT and GSIS experiments were determined by two-way ANOVA with post-hoc Tukey test. P-values for AUC analyses were determined by Student’s T-Test. *, p<0.05, **, p<0.01, ***<0.001, ns = not significant. All data are shown as mean ± SEM.

We collected blood at the 0- and 2-minute time points during the IP-GTT and assessed plasma insulin by ELISA (**Fig 3C, 3F**). Remarkably, induction of insulin secretion at 2 minutes was nearly absent in mutant mice. To ensure insulin sensitivity was not affected by *Steap2* ablation, we performed insulin tolerance tests (ITTs) (**Fig 3G, 3I**). No significant differences were observed in fold-change glucose (**Fig 3G, I**) or the AUC between control and mutant mice in either male or female cohorts (**Fig 3H**, **3J**), confirming a β-cell specific defect in glucose-stimulated insulin secretion.

### Continuous glucose monitoring of Steap2 mice by HD-XG telemetry probes

Under acute glucose challenge, *Steap2* mutant mice display an attenuated insulin secretory response. However, detecting responses during smaller glucose challenges, such as during the onset of feeding, is more challenging. This is due to the limited number of time points that can be sampled and the stress caused by mouse handling which may increase glucose levels (54; 55).

To address this issue, we performed continuous glucose monitoring using HD-XG telemetric probes in *Steap2* mice placed on an 18-week western diet (HFD) and housed at thermoneutrality (TN), hereafter referred to as “HFD”, to accelerate weight gain (**Fig 4A**). Weight gain between control and mutant mice were not significantly different in either male or female cohorts, but male control (slope = 1.40 g/week) and mutant mice (slope = 1.47 g/week) gained weight at a far greater pace than female control (slope = 0.82 g/week) and mutant mice (slope = 0.63 g/week) (**Fig S2A**).

**Figure 4:**
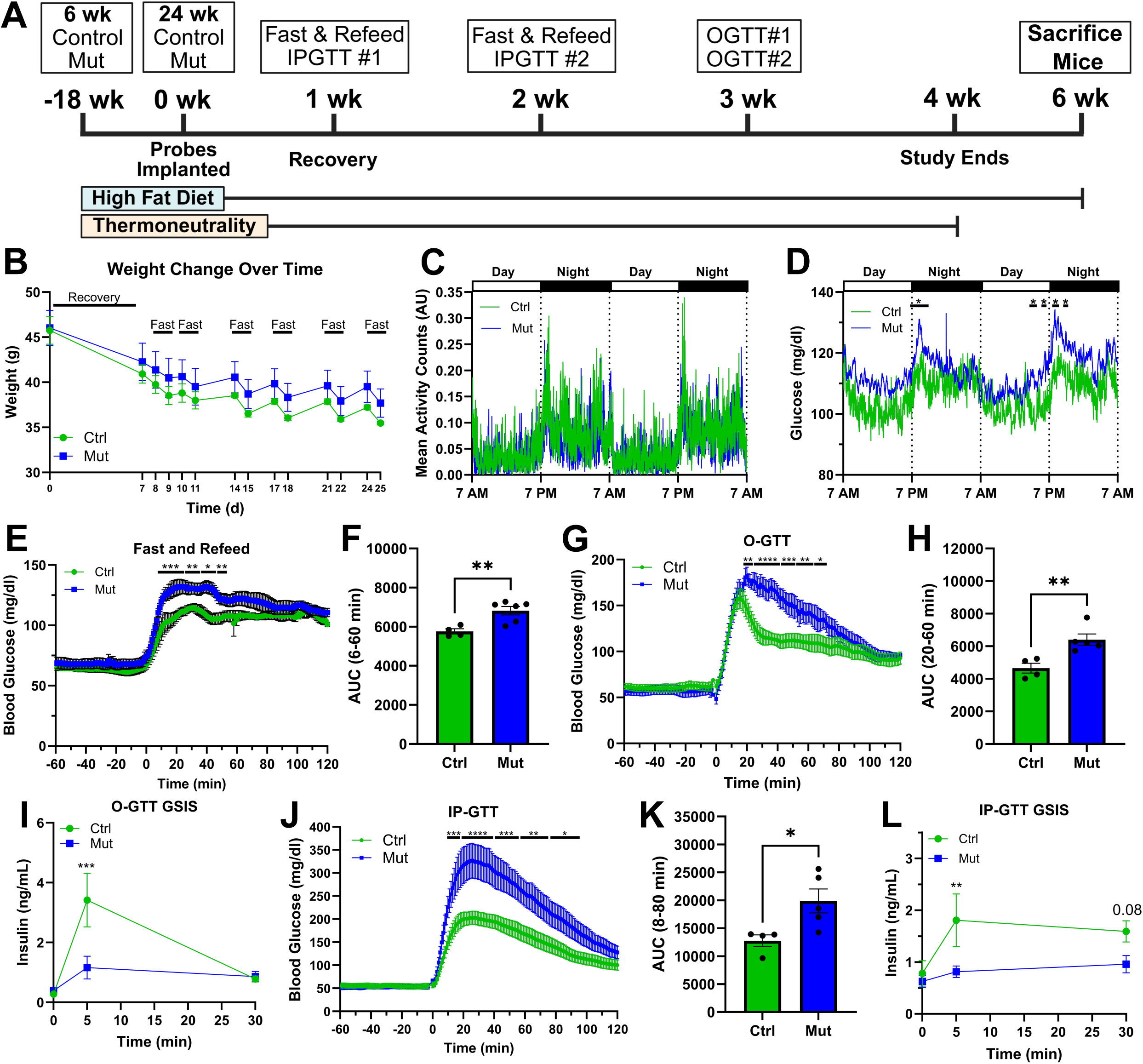
HD-XG telemetric continuous physiological monitoring of Western diet-fed *Steap2* mutant mice. **(A)** Experimental timeline. Male control (n = 5) and mutant mice (n = 6) aged 6 weeks were housed at thermoneutrality and fed a Western diet for 18-weeks to induce obesity. At 24-weeks of age, HD-XG probes were implanted and interventions conducted as indicated. **(B)** Weight curves. Weight decreased with surgical intervention and repeated fasts, though there were no significant differences between groups. **(C)** Mean activity counts (arbitrary units) of mice during day-night cycles in a 48-hour period, showing no significant differences between groups. Average of three 48-hour periods. **(D)** Random glucose over 48-hour period. Glucose is significantly more elevated in mutant mice during the beginning of the night cycle when mice first begin to feed. Average of three 48-hour periods. **(E)** 16-hour fast and 2-hour refeed of mice demonstrated significantly greater glucose excursions in mutant mice. Data are the average of two experiments. **(F)** AUC analysis of glucose excursion. **(G)** Oral glucose tolerance test (O-GTT) of control and mutant mice demonstrates a prolonged and greater glucose excursion in mutant mice. **(H)** AUC analysis of the O-GTT data. **(I)** *In vivo* GSIS during an O-GTT demonstrates significantly attenuated insulin secretion in mutant mice at 5 min, but not 30 min. **(J)** IP-GTT shows a prolonged and greater glucose excursion in mutant mice. **(K)** AUC analysis of the IP-GTT shows a greater glucose excursion in mutant mice. **(L)** *In vivo* GSIS during the IP-GTT demonstrates significantly attenuated insulin secretion in mutant mice at 5 min, but not 30 min. P-values for 48-hour measurements were determined by two-way ANOVA with post-hoc Holm test. P-Values for weight measurements, fast and refeed, IP-GTT, O-GTT, and GSIS were determined by two-way ANOVA with post-hoc Tukey Test. P-values for AUC analyses were determined by Student’s T-Test. *, p<0.05, **, p<0.01, ***<0.001, ns = not significant. Data shown as mean ± SEM.

Due to the limited weight gain in female mice, we opted to solely pursue probe implantation in male mice. After 18-weeks of HFD, male mice underwent surgical implantation of HD-XG probes and experiments outlined in **Figure 4A** were performed during the 3-week operational period of the glucose sensors. Although weight gradually decreased with the repeated fasts preceding GTTs and fast and refeed experiments, there was no difference by genotype **(Fig 4B)**. Mean activity accounts were not significantly different between groups during the 48-hour recovery period, and activity was decreased during the day and elevated during the night consistent with the nocturnal sleep patterns of mice **(Fig 4C)**. Blood glucose was elevated during the night phase and decreased during the day phase as expected from the feeding behavior of mice **(Fig 4D)**. Moreover, glucose dramatically rose during the transition period between day and night phases, corresponding to the onset of feeding, and blood glucose was significantly greater mutant mice than in controls.

Next, we assessed the glucose tolerance of HFD mice during 5-hour fast 2-hour refeed (F&R), IP-GTT and Oral (O-)GTT experiments. During the fasting/refeeding study, the glucose egress and AUC were significantly greater in mutant mice during the first hour of the refeed phase **(Fig 4E-F)**. Glucose was not significantly different 1-hour prior to the intervention during the fasting period. Oral glucose challenge with 1g D-glucose/kg bodyweight resulted in a significantly larger magnitude and duration of glucose egress in mutant mice **(Fig 4G-H)**. Tail vein blood was collected during a separate O-GTT at 0-, 5- and 30-min post glucose bolus, and insulin levels were measured. Plasma insulin was dramatically decreased in mutant mice compared to controls at the 5-min timepoint, but it was otherwise not significantly different at the 0 or 30 min timepoints **(Fig 4I)**. Glucose challenge with 1g D-glucose/kg bodyweight during the IP-GTT resulted in a dramatically greater magnitude and duration of glucose egress and AUC in mutant mice **(Fig 4J-K)**. Likewise, plasma insulin measured at 0-, 5- and 30-min timepoints from a separate IP-GTT was significantly decreased in mutant mice at 5-min, but not 0- or 30-min **(Fig 4L)**. Compared to the O-GTT, glucose egress was greater and insulin secretion was less marked during the IP-GTT in both control and mutant mice. Collectively, these findings indicate that the first phase of insulin secretion is blunted in mutant mice, while second phase insulin secretion as well as the incretin effect remain intact and are unaffected by *Steap2* deficiency. Interestingly, when comparing the glucose excursions during GTTs of mice subjected or not to tail vein bleed, there was a trending (p = 0.12-25) increase in the AUC in the handled mutant (B;mut) mice compared to unhandled mutant mice (NB;Mut) (**Fig S3C, F**).

### Ca^2+^ mobilization and dynamic insulin secretion are blunted in Steap2 deficient islets

Next, we investigated *in vitro* Ca^2+^ mobilization and insulin secretion dynamics. Isolated islets were loaded with the Fura-2 AM Ca^2+^ dye indicator and perfused with amino acids, 16.7 mM Glucose, 25 mM glucose and then KCl to induce maximal Ca^2+^ mobilization. Intracellular Ca^2+^ was measured every 10s over the course of the secretagogue ramp. Ca^2+^ mobilization was significantly blunted in male mutant mice during the glucose challenge phase (G 16.7-G 25 mM) as well as with KCl treatment **(Fig 5A).**

**Figure 5:**
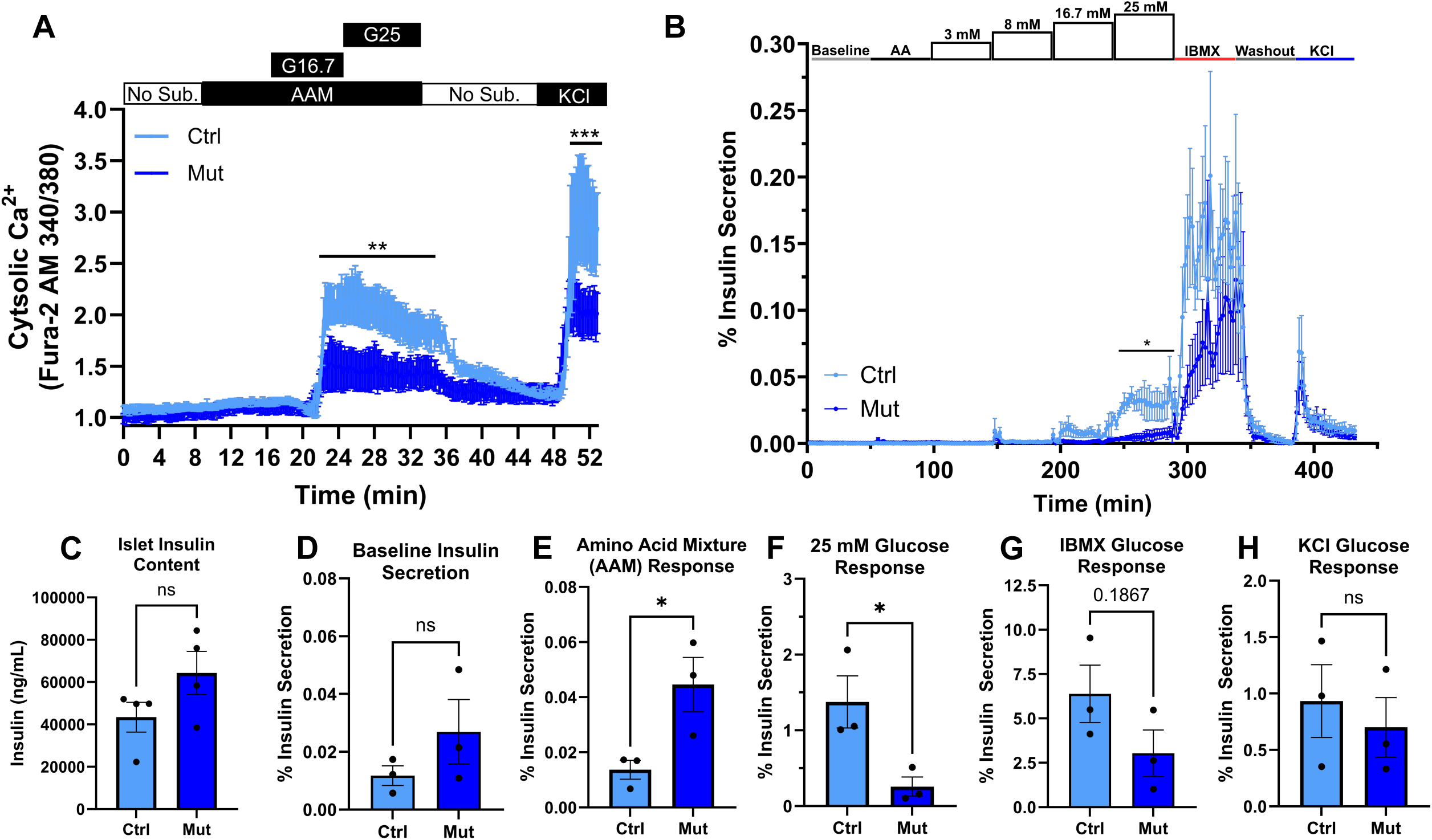
Glucose stimulated Ca^2+^ mobilization and insulin secretion is attenuated in isolated islets in *Steap2* mutant mice. **(A)** Intracellular Ca^2+^ response in islets isolated from 12-18-week-old male control (n = 5) and mutant (n = 3) mice perfused with no substrate (Krebs Ringer Bicarbonate Buffer), 16.7 mM Glucose, 25 mM glucose, no substrate washout and 30 mM KCl during which intracellular Ca^2+^ was measured by dual-wave fluorescence. Glucose and KCl-induced Ca^2+^ responses are significantly attenuated in mutant mice. **(B)** Insulin secretion from islets isolated from 12-18-week-old male control (n = 3) and mutant (n = 3) mice perfused with increasing glucose concentration in the presence of a physiological amino acid mixture (AAM) and then subsequently with IBMX and KCl. **(C)** ELISA measuring the total insulin content of perifused islets. **(D)** Area under the curve analysis of insulin secretion at baseline. **(E)** AUC analysis demonstrates a significantly elevated insulin secretion response to AAM in mutant mice. **(F)** AUC analysis demonstrates an attenuated insulin secretion response to 25 mM glucose in mutant mice. **(G-H)** AUC analysis of insulin response to IBMX and KCl. P-Values for intracellular Ca^2+^ and islet perifusion experiments were determined by two-way ANOVA with post-hoc Tukey Test. P-values for AUC analyses were determined by Student’s T-Test. *, p<0.05, **, p<0.01, ***<0.001, ns = not significant. All data are shown as mean ± SEM.

Next, we analyzed insulin secretion by islet perifusion. We challenged isolated islets with a secretagogue sequence comprised of amino acids with increasing glucose challenge (3 mM, 8 mM, 16.7 mM, 25 mM), then potentiated secretion with the cAMP activator 3-isobutyl-1-methylxanthine (IBMX, 0.1 mM) before depolarization with KCl **(Fig 5B)**. Insulin content was not significantly different between control and mutant islets **(Fig 5C)**. Baseline insulin secretion was not significantly different between mutant and control islet, but the response to physiological amino acid levels was significantly greater in mutant mice (**Fig 5D**). The insulin secretory response was significantly blunted in mutant mice during the 25 mM glucose challenge and was trending lower at G16.7 mM (p = 0.10-15) **(Fig 5B, 5E)**. The response to IBMX or KCl did not differ significantly by genotype (p = 0.187) (**Fig 5F,G)**. Taken together, *Steap2* mutant mice display impaired Ca^2+^ mobilization and insulin secretion in response to glucose stimulation, while cAMP elevating agents, such as IBMX and incretins, are able to rescue the insulin-secretory defect, as PKA and EPAC potentiate Ca^2+^ mobilization from VDCC and intracellular stores (56; 57).

### Steap2 deficiency results in changes to the islet transcriptome

Finally, we characterized the transcriptional adaptations of islets as a result of *Steap2* deficiency. We performed bulk RNA sequencing on RNA extracted from the isolated islets of male control (n = 3), male mutant (n = 3), female control (n = 4) and mutant (n = 2) age-matched, 8- to 9-week-old mice. Principal component analysis of RNA-seq dataset identified genotype (42%) and sex (31%) as major components accounting for overall sample variation **(Fig 6A).** Differential Gene Expression (DGE) analysis identified 439 upregulated and 198 down-regulated genes, respectively. The most differentially expressed (DE) genes are highlighted in the volcano plot in **Figure 6B**, and the expression of the 637 DE genes passing the above thresholds across genotype and sex is presented as a heatmap in **Figure 6C**.

**Figure 6:**
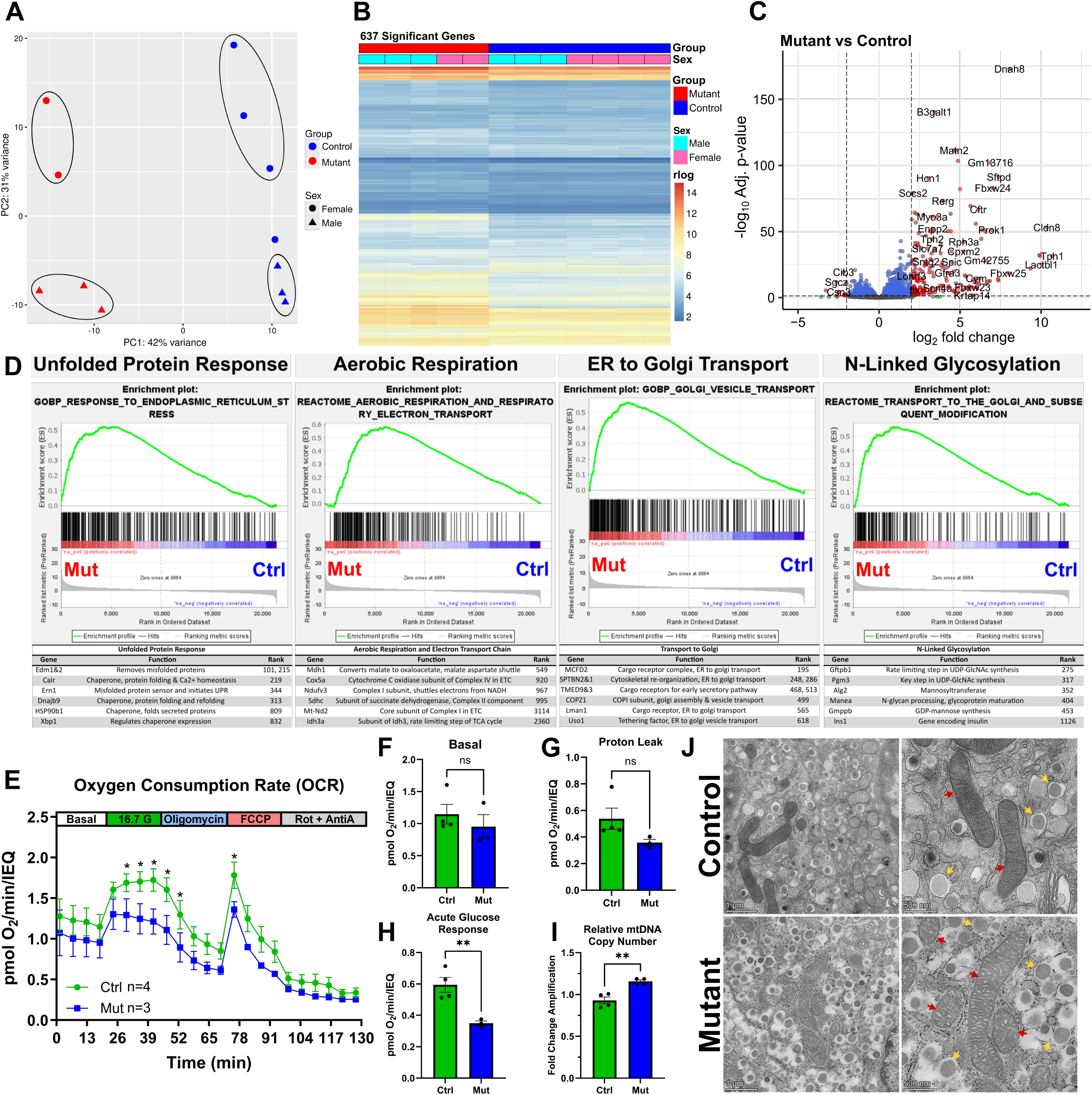
Transcriptome adaptations in *Steap2* mutant islets. **(A)** Principal component analysis plot of RNA-sequenced 8–9-week-old, male control (n = 3), male mutant (n = 3), female control (n = 4) and female mutant (n = 2) isolated islets showing that *Steap2* mutation primarily drives separation and clustering of samples. The plot was generated using DESeq2 plotPCA on regularized log transformed data (rlog) with blind=TRUE. **(B)** Heatmap showing the 637 differentially expressed genes with adjusted p-value<0.05 and |log_2_FoldChange| ≥ 1. The color scale corresponds to the rlog data with blind=FALSE. The blue and green bars indicate mutant and control genotypes, respectively, and the individual light blue and light pink bars indicate individual male and female mice, respectively. **(C)** Volcano plot of DESeq2 gene expression comparing control (n = 7) and mutant (n = 5) mouse islets. The color of each dot indicates whether the corresponding gene satisfies the thresholds of adjusted p-value<0.05 and |log_2_FoldChange| ≥1. **(D)** GSEA Enrichment plots for four pathways enriched in mutant mice are shown with the relative gene positions indicated by the straight lines (line plot) under each graph. Lines clustered to the left represent the “leading edge” and are the higher ranked genes in the pathway (according to the DESeq2 statistic). Genes of note within the leading edge are listed alongside their summarized function and rank in the pathway. **(E)** Oxygen consumption rate (OCR) (pmol O_2_/min/IEQ) of isolated islets normalized to islet equivalents (IEQs) from 8-9-week-old, male control (n = 4) and mutant (n = 3) mice in response to glucose (16.7 mM), oligomycin (5 μM), FCCP (2 μM), and rotenone/antimycin A (5 μM). **(F)** OCR at basal conditions (1 mM glucose, 0.5% FBS) **(G)** Seahorse segment analysis of OCR of proton leak (min oligomycin – min rotenone/AA) **(H)** Segment analysis of OCR (pmol/min/IEQ) in response to 16.7 mM glucose (max G16.7 – min Basal). **(I)** Genomic qPCR assessing fold-change difference in mitochondrial DNA (mtDNA) copy number between isolated islets from control (n = 4) and mutant mice (n = 4). **(J)** Transmission electron microscopy of pancreas sections at 11,000x and 22,000x magnification from 8-9-week-old, male, control (n = 2) and mutant (n = 2) mice. Red arrows denote mitochondria, yellow arrows denote insulin granules, and the white bars at the bottom left of each panel indicate the scale. Note that the cristae of mitochondria from mutant mice are swollen. P-values for islet seahorse were assessed by two-way ANOVA with post-hoc Tukey Test. P-values for seahorse segment analyses and qPCR were determined by Student’s T-Test. *, p<0.05, **, p<0.01, ***<0.001, ns = not significant. All data are shown as mean ± SEM.

Gene Set Enrichment Analysis (GSEA) (44) identified pathways related to the unfolded protein response, aerobic respiration and the electron transport chain, ER-derived protein trafficking to the Golgi, and N-linked glycosylation as upregulated in mutant mice **(Fig 6D)**. Given the enrichment of mitochondrial oxidative phosphorylation (OXPHOS) genes within the aerobic respiration pathway in *Steap2* mutant mice, we assessed whether mitochondrial function and morphology were affected. Interestingly, islet respirometry demonstrated that the oxygen consumption response (OCR) was instead significantly decreased in islets from mutant mice when stimulated with 16.7 mM glucose and 2 µM FCCP **(Fig 6E, H)**. While the acute glucose OCR was significantly lower in mutant islets, basal respiration and overall protein leak were not significantly different **(Fig 6F, G).** Moreover, qPCR for the mitochondrial gene *MT-RNR2* indicated that relative mitochondrial copy number was instead elevated in mutant mice **(Fig 6I)**. Yet, transmission electron microscopy imaging revealed that mitochondrial cristae of mutant mice (n = 2) were often swollen **(Fig 6J)**. These findings suggest that altered mitochondrial function during elevated metabolic flux, likely due to compromised ion balance affecting mitochondrial architecture, is impairing the production of ATP during glucose stimulation, thereby decreasing Ca^2+^ mobilization and subsequent insulin secretion.

TEM imaging also indicated that mutant mice (n = 4) exhibited far more empty insulin granules that were often aberrantly fused together than controls (n = 4) **(Fig S4A)**. Initially, it appeared that insulin folding and maturation may be altered and thus reflect the greater enrichment of ER stress, N-linked glycosylation and Golgi trafficking pathways in the GSEA data **(Fig 6D)**. However, even after accounting for differences in total islet protein **(Fig S4B)**, proinsulin and mature insulin were not significantly different between groups, and the PI:I ratio, although significantly different, was only lower by 0.5% in mutant mice **(Fig S4C-E)**. Thus, *Steap2* deficiency impairing mitochondrial respiration is likely the cause for the blunted GSIS observed in mutant mice.

## Discussion

*STEAP2* is a risk locus associated with altered glycemic control and type 2 diabetes risk (58; 59). Conditional deletion of *Steap2* in β-cells induced mitochondrial dysfunction impairing mitochondrial respiration in response to glucose challenge, resulting in blunted glucose-induced Ca^2+^ mobilization and insulin secretion.

In contrast to other mouse models achieving iron deprivation through embryonic deletion of IRP2 and TFRC1 (28; 29), *Steap2* depletion had no effect on random fasting glucose, weight gain or islet morphology. Instead, we observed glucose intolerance as early as 6 weeks during IP-GTT in conjunction with impaired in vivo GSIS during the first phase of insulin secretion. Interestingly, the difference in glucose excursion during IP-GTT between control and mutant mice was greatest in female mice. It is possible that this reflects recent observations that islets from female mice on C57bl6/j background have reduced peak insulin secretion and lower amplitude Ca^2+^ oscillations than those from male mice is when challenged with high glucose concentrations (60).

We employed state-of-the-art telemetric continuous glucose monitoring probes to enable high resolution characterization of mouse physiology and clarify the specific effect of *Steap2* deletion on glycemic control (34; 61). During the experimental period, we identified periods of greater glucose intolerance in mutant mice during the transition from the day to night phase, corresponding to feeding, as well as a stark difference in glucose excursion during the refeeding phase of fast and refeed experiments. Mutant mice did not otherwise exhibit differences in activity, random fed glucose or 16-h fasted glucose. Continuous glucose monitoring during IP-GTT and O-GTT recapitulated greater glucose intolerance in mutant mice and blunted insulin secretion during first phase insulin secretion. As expected, insulin secretion was markedly greater during O-GTT with a smaller glucose excursion in both control and mutant mice. Collectively, this suggests that *Steap2* deletion impairs first phase insulin secretion but neither the second phase insulin secretion nor the incretin effect (57), findings supported by islet perifusion and Ca^2+^ imaging studies.

RNA-seq analysis highlighted enrichment of pathways involved in the unfolded protein response, ER to Golgi transport and aerobic respiration. Specifically, genes involved in components of the electron transport chain were most enriched. These findings were corroborated by an increase in mtDNA copy number in isolated islets reflective of mitochondrial biogenesis. However, islet respirometry showed markedly decreased glucose-stimulated oxygen consumption in mutant mice in absence of altered basal respiration or proton leak. Interestingly, TEM showed that mitochondrial cristae of mutant mice were frequently swollen in a manner similar to the progressive mitochondrial dysfunction observed in TDP-43 associated neurodegenerative disease (62). Because we had observed both defects in GSIS as well as in mitochondrial function as determined by oxygen consumption experiments, we interpret the transcriptomic changes as an, ultimately insufficient, attempt of the mutant β-cells to compensate for impaired mitochondrial function through biogenesis.

In summary, we have established *Steap2* as the primary Steap metalloreductase in the β-cell that is indispensable optimal mitochondrial function and for the triggering pathway of first phase insulin secretion, suggesting that the impact of the *STEAP2* risk allele on glycemic traits is mediated by its function in β-cells.

## Acknowledgements

We thank the UPenn Transgenic & Chimeric Mouse Facility (RRID: SCR_022388) for their assistance with deriving the *Steap2^fl/fl^* allele, the UPenn Molecular Pathology and Imaging Core (RRID: SCR_022420) for with FFPE tissue sectioning and the UPenn Electron Microscopy Resource Lab (RRID: SCR_022375) for technical support with transmission electron microscopy. The authors thank BioRender (BioRender.com) for their considerable library of science related image assets. Lastly, we thank to the Rodent Metabolic Phenotyping and Islet Biology Cores of the UPenn Diabetes Research Center (RRID: SCR_022427; DK019525), for performing the continuous glucose monitoring experiments and islet studies.

## Funding

This work was supported by the NIH grants 3U01DK123594-02S2 to K.H.K and T32GM154643 to A.H.

## Duality of Interest

No conflicts of interest relevant to this publication were reported.

## Author Contributions

A.H., E.L.W, and K.H.K contributed to the study conceptualization and methodology. A.H. and K.H.K contributed to the project administration and writing of the original manuscript. A.H., E.L.W, M.T., K.K., Y.E., H.E., S.D., A.R., J.R., N.D., D.S. and E.M. contributed to the methodology, investigation, resources, and formal analysis. All authors contributed to reviewing and editing of the manuscript. K.H.K edited and finalized the manuscript and funding acquisition. K.H.K is the guarantor of this work and, as such, has full access to all the data in the study and takes responsibility for the integrity of the data and the accuracy of the data analysis.

## Notes

### Competing Interest Statement

The authors have declared no competing interest.

## References

1. The L: Diabetes: a defining disease of the 21st century. In The Lancet, 2023

2. Wilson PWF, Meigs JB, Sullivan L, Fox CS, Nathan DM, D’Agostino RB: Prediction of incident diabetes mellitus in middle-aged adults: The framingham offspring study. Archives of Internal Medicine 2007;167

3. Bhupathiraju SN, Hu FB: Epidemiology of obesity and diabetes and their cardiovascular complications. Circulation Research 2016;118

4. Abdullah A, Peeters A, de Courten M, Stoelwinder J: The magnitude of association between overweight and obesity and the risk of diabetes: A meta-analysis of prospective cohort studies. Diabetes Research and Clinical Practice 2010;89

5. Bell JA, Kivimaki M, Hamer M: Metabolically healthy obesity and risk of incident type 2 diabetes: A meta-analysis of prospective cohort studies. Obesity Reviews 2014;15

6. Fonseca VA: Defining and characterizing the progression of type 2 diabetes. Diabetes care 2009;32 Suppl 2

7. Goldfine AB, Bouche C, Parker RA, Kim C, Kerivan A, Soeldner JS, Martin BC, Warram JH, Kahn CR: Insulin resistance is a poor predictor of type 2 diabetes in individuals with no family history of disease. Proceedings of the National Academy of Sciences of the United States of America 2003;100

8. Vaag A, Henriksen JE, Beck-Nielsen H: Decreased insulin activation of glycogen synthase in skeletal muscles in young nonobese caucasian first-degree relatives of patients with non-insulin-dependent diabetes mellitus. Journal of Clinical Investigation 1992;89

9. Hollenbeck C, Reaven GM: Variations in Insulin-Stimulated Glucose Uptake in Healthy Individuals with Normal Glucose Tolerance. Journal of Clinical Endocrinology and Metabolism 1987;64

10. Willemsen G, Ward KJ, Bell CG, Christensen K, Bowden J, Dalgård C, Harris JR, Kaprio J, Lyle R, Magnusson PKE, Mather KA, Ordoňana JR, Perez-Riquelme F, Pedersen NL, Pietiläinen KH, Sachdev PS, Boomsma DI, Spector T: The Concordance and Heritability of Type 2 Diabetes in 34,166 Twin Pairs From International Twin Registers: The Discordant Twin (DISCOTWIN) Consortium. Twin Research and Human Genetics 2015;18

11. Avery AR, Duncan GE: Heritability of type 2 diabetes in the Washington state twin registry. Twin Research and Human Genetics 2019;22

12. Flannick J, Beer NL, Bick AG, Agarwala V, Molnes J, Gupta N, Burtt NP, Florez JC, Meigs JB, Taylor H, Lyssenko V, Irgens H, Fox E, Burslem F, Johansson S, Brosnan MJ, Trimmer JK, Newton-Cheh C, Tuomi T, Molven A, Wilson JG, O’Donnell CJ, Kathiresan S, Hirschhorn JN, Njølstad PR, Rolph T, Seidman JG, Gabriel S, Cox DR, Seidman CE, Groop L, Altshuler D: Assessing the phenotypic effects in the general population of rare variants in genes for a dominant Mendelian form of diabetes. In Nature Genetics, 2013

13. Suzuki K, Hatzikotoulas K, Southam L, Taylor HJ, Yin X, Lorenz KM, Mandla R, Huerta-Chagoya A, Melloni GEM, Kanoni S, Rayner NW, Bocher O, Arruda AL, Sonehara K, Namba S, Lee SSK, Preuss MH, Petty LE, Schroeder P, Vanderwerff B, Kals M, Bragg F, Lin K, Guo X, Zhang W, Yao J, Kim YJ, Graff M, Takeuchi F, Nano J, Lamri A, Nakatochi M, Moon S, Scott RA, Cook JP, Lee JJ, Pan I, Taliun D, Parra EJ, Chai JF, Bielak LF, Tabara Y, Hai Y, Thorleifsson G, Grarup N, Sofer T, Wuttke M, Sarnowski C, Gieger C, Nousome D, Trompet S, Kwak SH, Long J, Sun M, Tong L, Chen WM, Nongmaithem SS, Noordam R, Lim VJY, Tam CHT, Joo YY, Chen CH, Raffield LM, Prins BP, Nicolas A, Yanek LR, Chen G, Brody JA, Kabagambe E, An P, Xiang AH, Choi HS, Cade BE, Tan J, Broadaway KA, Williamson A, Kamali Z, Cui J, Thangam M, Adair LS, Adeyemo A, Aguilar-Salinas CA, Ahluwalia TS, Anand SS, Bertoni A, Bork-Jensen J, Brandslund I, Buchanan TA, Burant CF, Butterworth AS, Canouil M, Chan JCN, Chang LC, Chee ML, Chen J, Chen SH, Chen YT, Chen Z, Chuang LM, Cushman M, Danesh J, Das SK, de Silva HJ, Dedoussis G, Dimitrov L, Doumatey AP, Du S, Duan Q, Eckardt KU, Emery LS, Evans DS, Evans MK, Fischer K, Floyd JS, Ford I, Franco OH, Frayling TM, Freedman BI, Genter P, Gerstein HC, Giedraitis V, González-Villalpando C, González-Villalpando ME, Gordon-Larsen P, Gross M, Guare LA, Hackinger S, Hakaste L, Han S, Hattersley AT, Herder C, Horikoshi M, Howard AG, Hsueh W, Huang M, Huang W, Hung YJ, Hwang MY, Hwu CM, Ichihara S, Ikram MA, Ingelsson M, Islam MT, Isono M, Jang HM, Jasmine F, Jiang G, Jonas JB, Jørgensen T, Kamanu FK, Kandeel FR, Kasturiratne A, Katsuya T, Kaur V, Kawaguchi T, Keaton JM, Kho AN, Khor CC, Kibriya MG, Kim DH, Kronenberg F, Kuusisto J, Läll K, Lange LA, Lee KM, Lee MS, Lee NR, Leong A, Li L, Li Y, Li-Gao R, Ligthart S, Lindgren CM, Linneberg A, Liu CT, Liu J, Locke AE, Louie T, Luan Ja, Luk AO, Luo X, Lv J, Lynch JA, Lyssenko V, Maeda S, Mamakou V, Mansuri SR, Matsuda K, Meitinger T, Melander O, Metspalu A, Mo H, Morris AD, Moura FA, Nadler JL, Nalls MA, Nayak U, Ntalla I, Okada Y, Orozco L, Patel SR, Patil S, Pei P, Pereira MA, Peters A, Pirie FJ, Polikowsky HG, Porneala B, Prasad G, Rasmussen-Torvik LJ, Reiner AP, Roden M, Rohde R, Roll K, Sabanayagam C, Sandow K, Sankareswaran A, Sattar N, Schönherr S, Shahriar M, Shen B, Shi J, Shin DM, Shojima N, Smith JA, So WY, Stančáková A, Steinthorsdottir V, Stilp AM, Strauch K, Taylor KD, Thorand B, Thorsteinsdottir U, Tomlinson B, Tran TC, Tsai FJ, Tuomilehto J, Tusie-Luna T, Udler MS, Valladares-Salgado A, van Dam RM, van Klinken JB, Varma R, Wacher-Rodarte N, Wheeler E, Wickremasinghe AR, van Dijk KW, Witte DR, Yajnik CS, Yamamoto K, Yamamoto K, Yoon K, Yu C, Yuan JM, Yusuf S, Zawistowski M, Zhang L, Zheng W, Kanona S, van Heel DA, Raffel LJ, Igase M, Ipp E, Redline S, Cho YS, Lind L, Province MA, Fornage M, Hanis CL, Ingelsson E, Zonderman AB, Psaty BM, Wang YX, Rotimi CN, Becker DM, Matsuda F, Liu Y, Yokota M, Kardia SLR, Peyser PA, Pankow JS, Engert JC, Bonnefond A, Froguel P, Wilson JG, Sheu WHH, Wu JY, Hayes MG, Ma RCW, Wong TY, Mook-Kanamori DO, Tuomi T, Chandak GR, Collins FS, Bharadwaj D, Paré G, Sale MM, Ahsan H, Motala AA, Shu XO, Park KS, Jukema JW, Cruz M, Chen YDI, Rich SS, McKean-Cowdin R, Grallert H, Cheng CY, Ghanbari M, Tai ES, Dupuis J, Kato N, Laakso M, Köttgen A, Koh WP, Bowden DW, Palmer CNA, Kooner JS, Kooperberg C, Liu S, North KE, Saleheen D, Hansen T, Pedersen O, Wareham NJ, Lee J, Kim BJ, Millwood IY, Walters RG, Stefansson K, Ahlqvist E, Goodarzi MO, Mohlke KL, Langenberg C, Haiman CA, Loos RJF, Florez JC, Rader DJ, Ritchie MD, Zöllner S, Mägi R, Marston NA, Ruff CT, van Heel DA, Finer S, Denny JC, Yamauchi T, Kadowaki T, Chambers JC, Ng MCY, Sim X, Below JE, Tsao PS, Chang KM, McCarthy MI, Meigs JB, Mahajan A, Spracklen CN, Mercader JM, Boehnke M, Rotter JI, Vujkovic M, Voight BF, Morris AP, Zeggini E: Genetic drivers of heterogeneity in type 2 diabetes pathophysiology. Nature 2024;627

14. Flannick J, Mercader JM, Fuchsberger C, Udler MS, Mahajan A, Wessel J, Teslovich TM, Caulkins L, Koesterer R, Barajas-Olmos F, Blackwell TW, Boerwinkle E, Brody JA, Centeno-Cruz F, Chen L, Chen S, Contreras-Cubas C, Córdova E, Correa A, Cortes M, DeFronzo RA, Dolan L, Drews KL, Elliott A, Floyd JS, Gabriel S, Garay-Sevilla ME, García-Ortiz H, Gross M, Han S, Heard-Costa NL, Jackson AU, Jørgensen ME, Kang HM, Kelsey M, Kim BJ, Koistinen HA, Kuusisto J, Leader JB, Linneberg A, Liu CT, Liu J, Lyssenko V, Manning AK, Marcketta A, Malacara-Hernandez JM, Martínez-Hernández A, Matsuo K, Mayer-Davis E, Mendoza-Caamal E, Mohlke KL, Morrison AC, Ndungu A, Ng MCY, O’Dushlaine C, Payne AJ, Pihoker C, Post WS, Preuss M, Psaty BM, Vasan RS, Rayner NW, Reiner AP, Revilla-Monsalve C, Robertson NR, Santoro N, Schurmann C, So WY, Soberón X, Stringham HM, Strom TM, Tam CHT, Thameem F, Tomlinson B, Torres JM, Tracy RP, van Dam RM, Vujkovic M, Wang S, Welch RP, Witte DR, Wong TY, Atzmon G, Barzilai N, Blangero J, Bonnycastle LL, Bowden DW, Chambers JC, Chan E, Cheng CY, Cho YS, Collins FS, de Vries PS, Duggirala R, Glaser B, Gonzalez C, Gonzalez ME, Groop L, Kooner JS, Kwak SH, Laakso M, Lehman DM, Nilsson P, Spector TD, Tai ES, Tuomi T, Tuomilehto J, Wilson JG, Aguilar-Salinas CA, Bottinger E, Burke B, Carey DJ, Chan JCN, Dupuis J, Frossard P, Heckbert SR, Hwang MY, Kim YJ, Kirchner HL, Lee JY, Lee J, Loos RJF, Ma RCW, Morris AD, O’Donnell CJ, Palmer CNA, Pankow J, Park KS, Rasheed A, Saleheen D, Sim X, Small KS, Teo YY, Haiman C, Hanis CL, Henderson BE, Orozco L, Tusié-Luna T, Dewey FE, Baras A, Gieger C, Meitinger T, Strauch K, Lange L, Grarup N, Hansen T, Pedersen O, Zeitler P, Dabelea D, Abecasis G, Bell GI, Cox NJ, Seielstad M, Sladek R, Meigs JB, Rich SS, Rotter JI, Altshuler D, Burtt NP, Scott LJ, Morris AP, Florez JC, McCarthy MI, Boehnke M: Exome sequencing of 20,791 cases of type 2 diabetes and 24,440 controls. Nature 2019;570

15. Costanzo MC, Akolkar B, Claussnitzer M, Florez JC, Gloyn AL, Grant SFA, Kaestner KH, Manning AK, Mohlke KL, Parker SCJ, Titchenell PM, Udler MS, Jones MA, Kamphaus TN, Fischer RA, McCarthy MI, Miller MR, Boehnke M, Flannick J, Burtt NP: Accelerating Medicines Partnership in Type 2 Diabetes and Common Metabolic Diseases: Collaborating to Maximize the Value of Genetic and Genomic Data. Diabetes 2025;74

16. Dornbos P, Singh P, Jang DK, Mahajan A, Biddinger SB, Rotter JI, McCarthy MI, Flannick J: Evaluating human genetic support for hypothesized metabolic disease genes. In Cell Metabolism, 2022

17. Ohgami RS, Campagna DR, McDonald A, Fleming MD: The Steap proteins are metalloreductases. Blood 2006;108

18. Cai Q, Jing C, Wang X, Xing X, Liu W: STEAP Proteins: Roles in disease biology and potential for therapeutic intervention. International Journal of Biological Macromolecules 2025;309:142797–142797

19. Ben Zichri-David S, Shkuri L, Ast T: Pulling back the mitochondria’s iron curtain. NPJ Metab Health Dis 2025;3:6

20. Ford ES, Cogswell ME: Diabetes and serum ferritin concentration among U.S. adults. Diabetes Care 1999;22

21. Jiang R, Manson JE, Meigs JB, Ma J, Rifai N, Hu FB: Body iron stores in relation to risk of type 2 diabetes in apparently healthy women. JAMA 2004;291:711–717

22. Hudson DM, Curtis SB, Smith VC, Griffiths TAM, Wong AYK, Scudamore CH, Buchan AMJ, MacGillivray RTA: Human hephaestin expression is not limited to enterocytes of the gastrointestinal tract but is also found in the antrum, the enteric nervous system, and pancreatic β-cells. American Journal of Physiology - Gastrointestinal and Liver Physiology 2010;298

23. Koch RO, Zoller H, Theurl I, Obrist P, Egg G, Strohmayer W, Vogel W, Weiss G: Distribution of DMT 1 within the human glandular system. Histology and Histopathology 2003;18

24. Lenzen S, Drinkgern J, Tiedge M: Low antioxidant enzyme gene expression in pancreatic islets compared with various other mouse tissues. Free Radical Biology and Medicine 1996;20

25. Wu K, Fei L, Wang X, Lei Y, Yu L, Xu W, Chen J, Zhu E, Zhong M, Huang M, Xi J, Yin F, Yan Z, Zhao X, Tang C, Patzak A, Liu X, Zheng Z: ZIP14 is involved in iron deposition and triggers ferroptosis in diabetic nephropathy. Metallomics 2022;14

26. Hardy AB, Prentice KJ, Froese S, Liu Y, Andrews GK, Wheeler MB: Zip4 mediated zinc influx stimulates insulin secretion in pancreatic beta cells. PLoS ONE 2015;10

27. Berthault C, Staels W, Scharfmann R: Purification of pancreatic endocrine subsets reveals increased iron metabolism in beta cells. Molecular Metabolism 2020;42

28. Van Mulders A, Willems L, Coenen S, Bourgeois S, Yi X, Tong Y, Leuckx G, Heremans Y, Pierreux J, Degroote L, Sawatani T, Vinci C, Ates G, Massie A, Carlotti F, de Koning E, Mandrup-Poulsen T, Zelinsky C, Goderis S, Ghesquière B, Scharfmann R, Cnop M, De Leu N, Staels W: Iron deficiency induces maturation-dependent loss of pancreatic β-cells. Nature communications 2026;17

29. Santos MCFd, Anderson CP, Neschen S, Zumbrennen-Bullough KB, Romney SJ, Kahle-Stephan M, Rathkolb B, Gailus-Durner V, Fuchs H, Wolf E, Rozman J, de Angelis MH, Cai WM, Rajan M, Hu J, Dedon PC, Leibold EA: Irp2 regulates insulin production through iron-mediated Cdkal1-catalyzed tRNA modification. Nature Communications 2020;11

30. Hansen JB, Tonnesen MF, Madsen AN, Hagedorn PH, Friberg J, Grunnet LG, Heller RS, Nielsen AO, Storling J, Baeyens L, Anker-Kitai L, Qvortrup K, Bouwens L, Efrat S, Aalund M, Andrews NC, Billestrup N, Karlsen AE, Holst B, Pociot F, Mandrup-Poulsen T: Divalent metal transporter 1 regulates iron-mediated Ros and pancreatic β cell fate in response to cytokines. Cell Metabolism 2012;16

31. Wellen KE, Fucho R, Gregor MF, Furuhashi M, Morgan C, Lindstad T, Vaillancourt E, Gorgun CZ, Saatcioglu F, Hotamisligil GS: Coordinated Regulation of Nutrient and Inflammatory Responses by STAMP2 Is Essential for Metabolic Homeostasis. Cell 2007;129

32. Lindstad T, Qu S, Sikkeland J, Jin Y, Kristian A, Mælandsmo GM, Collas P, Saatcioglu F: STAMP2 is required for human adipose-derived stem cell differentiation and adipocyte-facilitated prostate cancer growth in vivo. Oncotarget 2017;8

33. Herrera PL, Orci L, Vassalli JD. Two transgenic approaches to define the cell lineages in endocrine pancreas development. In Molecular and Cellular Endocrinology.

34. Korstanje R, Ryan JL, Savage HS, Lyons BL, Kane KG, Sukoff Rizzo SJ: Continuous Glucose Monitoring in Female NOD Mice Reveals Daily Rhythms and a Negative Correlation With Body Temperature. Endocrinology 2017;158:2707–2712

35. Doliba NM, Qin W, Najafi H, Liu C, Buettger CW, Sotiris J, Collins HW, Li C, Stanley CA, Wilson DF, Grimsby J, Sarabu R, Naji A, Matschinsky FM: Glucokinase activation repairs defective bioenergetics of islets of Langerhans isolated from type 2 diabetics. American Journal of Physiology - Endocrinology and Metabolism 2012;302

36. Chen KH, Doliba N, May CL, Roman J, Ustione A, Tembo T, Negron A, Radovick S, Piston DW, Glaser B, Kaestner KH, Matschinsky FM: Genetic activation of glucokinase in a minority of pancreatic beta cells causes hypoglycemia in mice. Life Sciences 2022;309

37. Swisa A, Kieckhaefer J, Daniel SG, El-Mekkoussi H, Kolev HM, Tigue M, Jin C, Assenmacher CA, Dohnalová L, Thaiss CA, Karlsson NG, Bittinger K, Kaestner KH: The evolutionarily ancient FOXA transcription factors shape the murine gut microbiome via control of epithelial glycosylation. Developmental Cell 2024;59

38. Rooney JP, Ryde IT, Sanders LH, Howlett EV, Colton MD, Germ KE, Mayer GD, Timothy Greenamyre J, Meyer JN: PCR based determination of mitochondrial DNA copy number in multiple species. Methods in Molecular Biology 2015;1241

39. Kin T: Islet isolation for clinical transplantation. In Advances in Experimental Medicine and Biology, 2010

40. Chen S, Zhou Y, Chen Y, Gu J. Fastp: An ultra-fast all-in-one FASTQ preprocessor. In *Bioinformatics*.

41. Dobin A, Davis CA, Schlesinger F, Drenkow J, Zaleski C, Jha S, Batut P, Chaisson M, Gingeras TR: STAR: Ultrafast universal RNA-seq aligner. Bioinformatics 2013;29

42. Liao Y, Smyth GK, Shi W: FeatureCounts: An efficient general purpose program for assigning sequence reads to genomic features. Bioinformatics 2014;30

43. Love MI, Huber W, Anders S: Moderated estimation of fold change and dispersion for RNA-seq data with DESeq2. Genome Biology 2014;15

44. Subramanian A, Tamayo P, Mootha VK, Mukherjee S, Ebert BL, Gillette MA, Paulovich A, Pomeroy SL, Golub TR, Lander ES, Mesirov JP: Gene set enrichment analysis: A knowledge-based approach for interpreting genome-wide expression profiles. Proceedings of the National Academy of Sciences of the United States of America 2005;102

45. Manduchi E, Descamps HC, Schug J, Da T, Lahori D, El-Mekkoussi H, Betts MR, Kaestner KH: No Evidence for Persistent Enteroviral B Infection of Pancreatic Islets in Patients With Type 1 Diabetes and Prediabetes From RNA Sequencing Data. Diabetes 2024;73

46. Kaestner KH, Powers AC, Naji A, Consortium H, Atkinson MA: NIH Initiative to Improve Understanding of the Pancreas, Islet, and Autoimmunity in Type 1 Diabetes: The Human Pancreas Analysis Program (HPAP). Diabetes 2019;68:1394–1402

47. Shapira SN, Naji A, Atkinson MA, Powers AC, Kaestner KH: Understanding islet dysfunction in type 2 diabetes through multidimensional pancreatic phenotyping: The Human Pancreas Analysis Program. In Cell Metabolism, 2022

48. Chen K, Wang L, Shen J, Tsai AL, Zhou M, Wu G: Mechanism of stepwise electron transfer in six-transmembrane epithelial antigen of the prostate (STEAP) 1 and 2. eLife 2023;12

49. Ahmad S, Jose da Costa Gonzales L, Bowler-Barnett EH, Rice DL, Kim M, Wijerathne S, Luciani A, Kandasaamy S, Luo J, Watkins X, Turner E, Martin MJ, UniProt C: The UniProt website API: facilitating programmatic access to protein knowledge. Nucleic Acids Res 2025;53:W547–W553

50. Estall JL, Screaton RA: Of mice and men, redux: Modern challenges in β cell gene targeting. In Endocrinology (United States), 2021

51. Lee JY, Ristow M, Lin X, White MF, Magnuson MA, Hennighausen L: RIP-Cre revisited, evidence for impairments of pancreatic β-cell function. Journal of Biological Chemistry 2006;281

52. Teitelman G, Kedees M: Mouse insulin cells expressing an inducible RIPCre transgene are functionally impaired. Journal of Biological Chemistry 2015;290

53. Carboneau BA, Le TDV, Dunn JC, Gannon M: Unexpected effects of the MIP-CreER transgene and tamoxifen on β-cell growth in C57Bl6/J male mice. Physiological Reports 2016;4

54. Ghosal S, Nunley A, Mahbod P, Lewis AG, Smith EP, Tong J, D’Alessio DA, Herman JP: Mouse handling limits the impact of stress on metabolic endpoints. Physiology and Behavior 2015;150

55. Sim YB, Park SH, Kang YJ, Kim SM, Lee JK, Jung JS, Suh HW: The regulation of blood glucose level in physical and emotional stress models: Possible involvement of adrenergic and glucocorticoid systems. Archives of Pharmacal Research 2010;33

56. Stožer A, Leitgeb EP, Pohorec V, Dolenšek J, Bombek LK, Gosak M, Klemen MS: The role of camp in beta cell stimulus–secretion and intercellular coupling. In Cells, 2021

57. Campbell JE, Newgard CB: Mechanisms controlling pancreatic islet cell function in insulin secretion. In Nature Reviews Molecular Cell Biology, 2021

58. Brotman SM, El-Sayed Moustafa JS, Guan L, Broadaway KA, Wang D, Jackson AU, Welch R, Currin KW, Tomlinson M, Vadlamudi S, Stringham HM, Roberts AL, Lakka TA, Oravilahti A, Fernandes Silva L, Narisu N, Erdos MR, Yan T, Bonnycastle LL, Raulerson CK, Raza Y, Yan X, Parker SCJ, Kuusisto J, Pajukanta P, Tuomilehto J, Collins FS, Boehnke M, Love MI, Koistinen HA, Laakso M, Mohlke KL, Small KS, Scott LJ: Adipose tissue eQTL meta-analysis highlights the contribution of allelic heterogeneity to gene expression regulation and cardiometabolic traits. Nature Genetics 2025;57

59. Su C, Gao L, May CL, Pippin JA, Boehm K, Lee M, Liu C, Pahl MC, Golson ML, Naji A, Grant SFA, Wells AD, Kaestner KH: 3D chromatin maps of the human pancreas reveal lineage-specific regulatory architecture of T2D risk. Cell Metabolism 2022;34

60. Jo S, Beetch M, Gustafson E, Wong A, Oribamise E, Chung G, Vadrevu S, Satin LS, Bernal-Mizrachi E, Alejandro EU: Sex Differences in Pancreatic β-Cell Physiology and Glucose Homeostasis in C57BL/6J Mice. Journal of the Endocrine Society 2023;7

61. Brockway R, Tiesma S, Bogie H, White K, Fine M, O’Farrell L, Michael M, Cox A, Coskun T: Fully implantable arterial blood glucose device for metabolic research applications in rats for two months. Journal of Diabetes Science and Technology 2015;9

62. Wang P, Deng J, Dong J, Liu J, Bigio EH, Mesulam M, Wang T, Sun L, Wang L, Lee AYL, McGee WA, Chen X, Fushimi K, Zhu L, Wu JY: TDP-43 induces mitochondrial damage and activates the mitochondrial unfolded protein response. PLoS Genetics 2019;15

