## Supplementary Materials for "The ortholog of the Human T2D Risk Gene *STEAP2* is Required for Glucose Stimulated Insulin Secretion in Mice"

Supplementary Materials – Hawthorne et al.

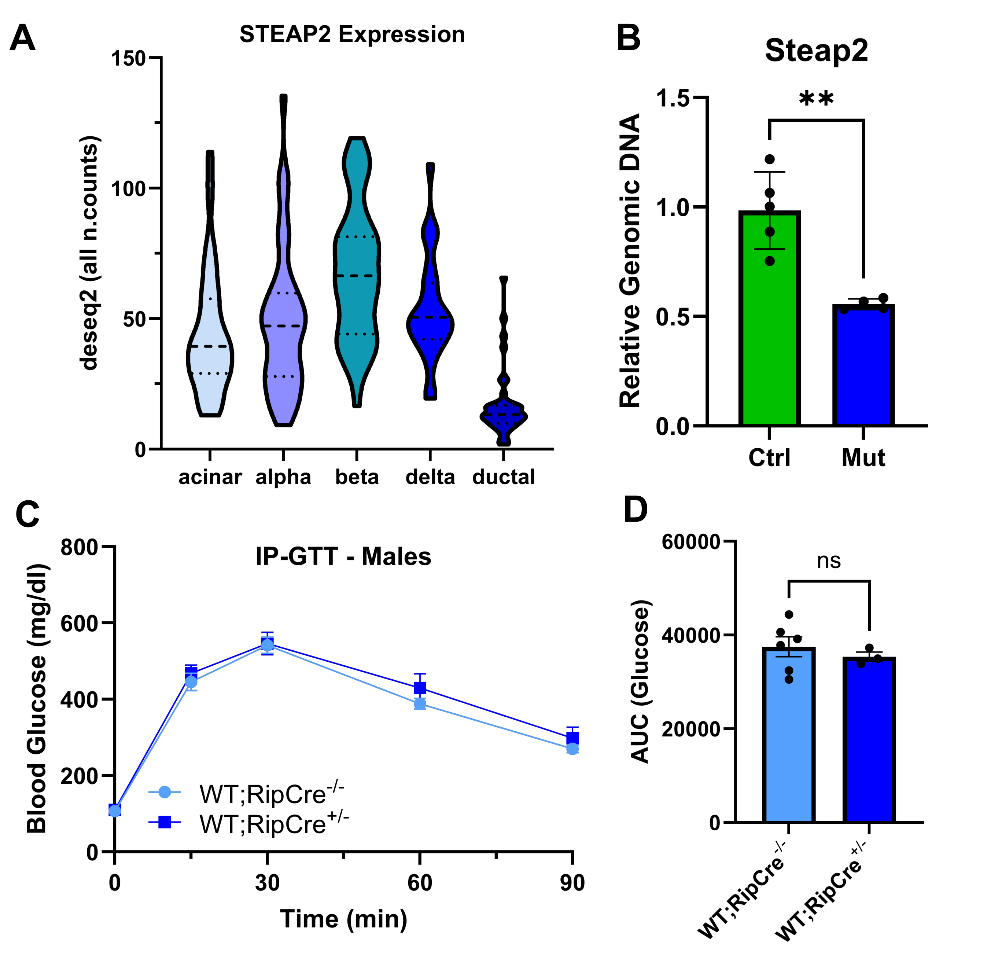

**Supplementary Figure 1: The RipCre allele does not confer a glucose intolerance phenotype.** **(A)** Violin plots displaying single-cell RNA Seq deseq2-normalized pseudobulk counts of *STEAP2* in human pancreatic cell types. Note that *STEAP2* counts are greatest in β-cells with the relative abundance of counts being β > δ > α > acinar > ductal. **(B)** Genomic DNA qPCR of *Steap2* exon 4 in isolated islets from 12-18 wk old mice (Control n=5, Mutant n=4) confirming ~50% reduction of *Steap2* DNA abundance. **(C)** No significant differences in blood glucose (mg/dl) between wild type (*WT;RipCre^-/-^*) (n = 6) and *WT;RipCre^+/-^* mice (n = 3) during 16-hour fasted IP-GTT were observed. **(D)** AUC analysis of IP-GTT demonstrates no significant differences in glucose egress. P-values for the IP-GTT were determined by two-way ANOVA with post-hoc Tukey test. P-values for AUC analyses and qPCR were determined by Student’s T-Test. *, p<0.05, **, p<0.01, ***<0.001, ns = not significant. All data are shown as mean ± SEM.

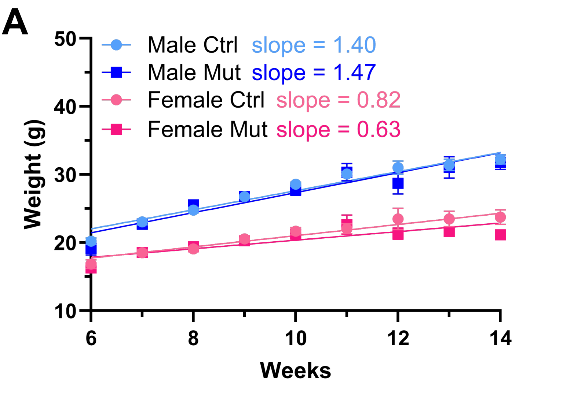

**Supplemental Figure 2: Male** ***Steap2* mutant mice on high fat diet (HFD) and housed at thermoneutrality (TN) gain weight at a greater rate than female mice.** **(A)** Weights and accompanying slope of weight gain over time (g/week) of male control (n = 8), male mutant (n = 6), female control (n = 5), and female mutant (n = 5) mice between 6-14 weeks of age. Note the considerably greater slope in male mice. P-values for weight gain over time were determined by two-way ANOVA with post-hoc Tukey test. *, p<0.05, **, p<0.01, ***<0.001, ns = not significant. All data are shown as mean ± SEM.

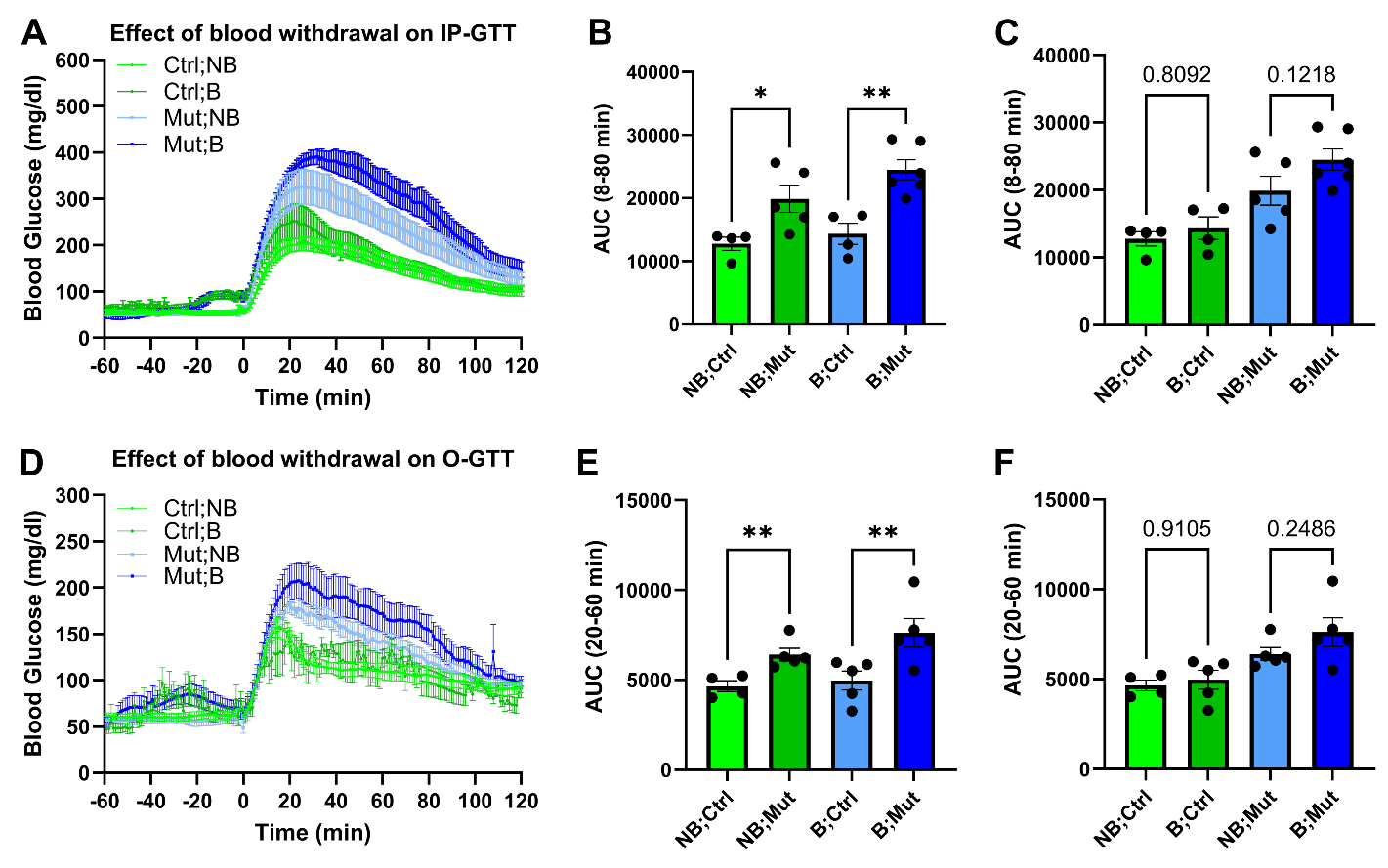

**Supplementary Figure 3: Blood withdrawals during GTTs result in increased glucose intolerance.** **(A)** IP-GTT of mice where tail vein blood was withdrawn from male control (n = 4) and mutant (n = 6) mice as well as was not (NB) withdrawn from male control (n = 4) and mutant (n = 5) mice fed a western diet for 18-weeks and housed at thermoneutrality (30°C). Multiple comparisons by two-way ANOVA did not reach significance at any timepoint when comparing NB;Ctrl to B;Ctrl or NB;Mut to B;Mut, but was trending (P-value = 0.058-0.15) between 5-120 min. **(B-C)** Area under the curve analysis comparing control and mutant mice across genotype and blood withdrawal status. **(D)** O-GTT of mice where tail vein blood was withdrawn (B) from male control (n = 4) and mutant (n = 5) mice as well as was not withdrawn (NB) from male control (n = 4) and mutant (n = 4) mice fed a western diet for 18-weeks and housed at thermoneutrality (30°C). **(E-F)** Area under the curve analysis comparing control and mutant mice across genotype and blood withdrawal status. P-values for the IP-GTT were determined by two-way ANOVA with post-hoc Tukey test. P-values for AUC analyses were determined by Student’s T-Test. *, p<0.05, **, p<0.01, ***<0.001, ns = not significant. All data are shown as mean ± SEM.

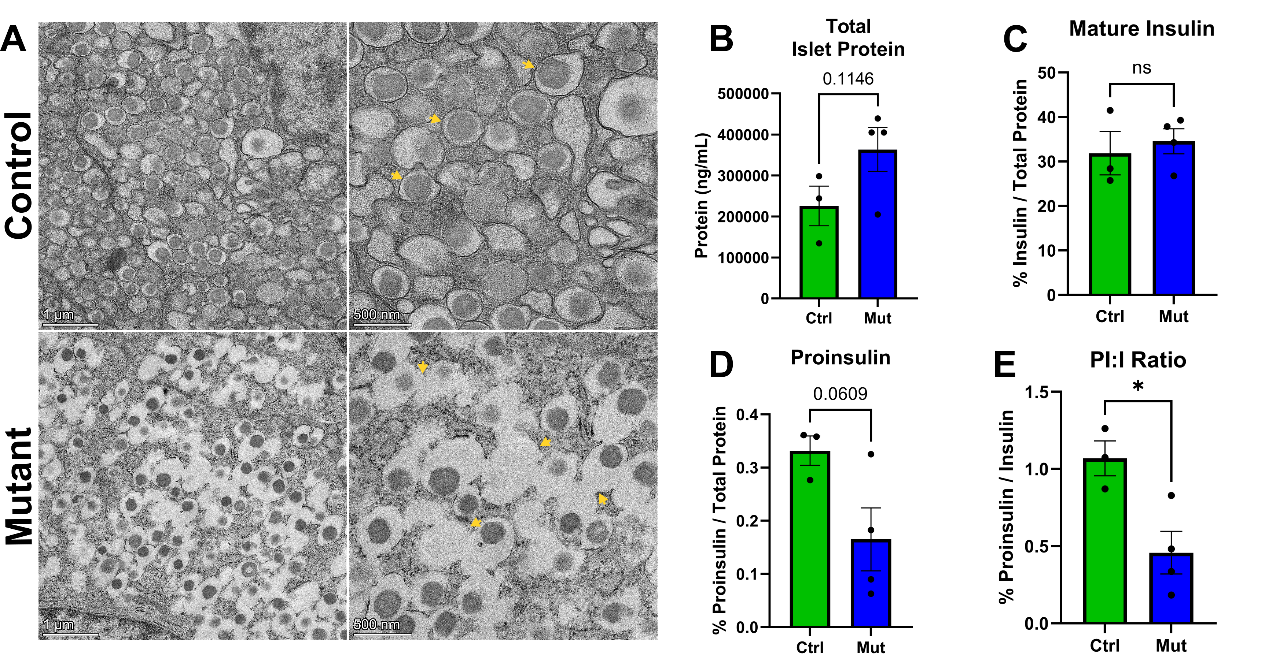

**Supplementary Figure 4: Empty and fused insulin granules are increased in *Steap2* mutant mice.** **(A)** Transmission electron microscopy of pancreas sections from 8-9-week-old, male, control (n = 4) and mutant (n = 4) mice. Yellow arrows highlight insulin granules. Note that the number of empty and aberrantly fused insulin granules are increased in mutant mice. **(B)** Bradford assay assessing total islet protein concentration from isolated islet lysates **(C)** ELISA measuring mature insulin and **(D)** proinsulin as percent of total protein content from isolated islet lysates of 12-18-week-old male, control (n = 3) and mutant (n = 4) mice. **(E)** Ratio of proinsulin to insulin in isolated islets. P-values for ELISA were determined by Student’s T-Test. *, p<0.05, **, p<0.01, ***<0.001, ns = not significant. All data are shown as mean ± SEM.

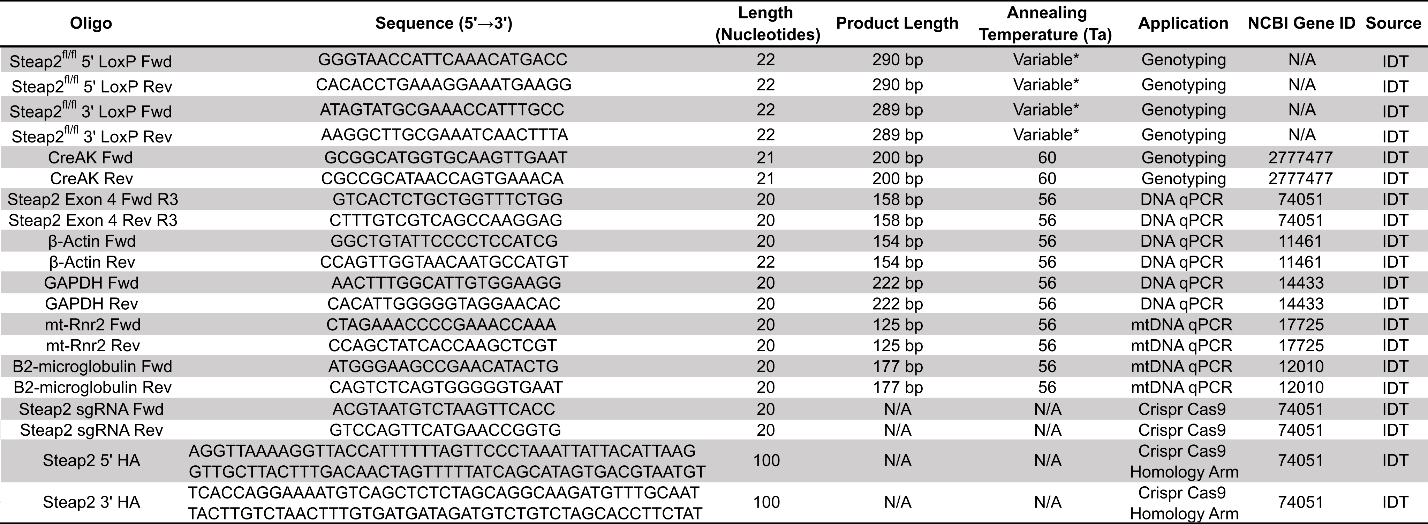
**Supplementary Table 1: Primers and Crispr-Cas9 guide RNAs and homology arm sequences used.** List of primer pairs utilized in genotyping, qPCR as well as oligos used in Crispr-Ca9 mediated derivation of *Steap2^fl/fl^* mouse line. Oligo names, sequences (5’🡪3’), annealing temperatures (*T_a_*), expected amplicon sizes (bp), NCBI gene IDs, and manufacturer are listed.

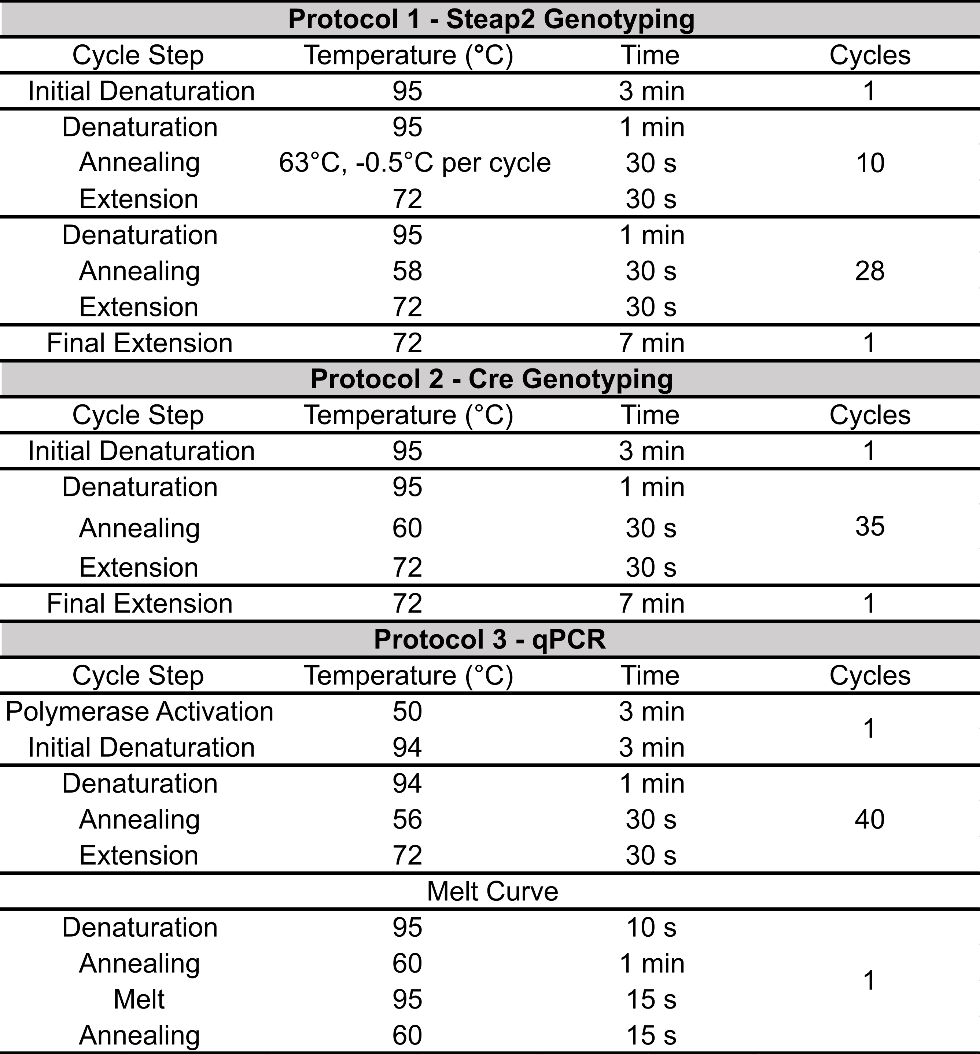

**Supplementary Table 2: PCR cycling conditions used.** List of PCR protocols and their respective cycling conditions employed to genotype the mice for *Steap2^fl/fl^* allele and the Herrera *RipCre* allele as well to perform Sybr Green genomic qPCR. The protocols highlighted in grey as well as respective cycle steps, temperatures of each step, duration (time) of each step as well as total cycles per step are listed.

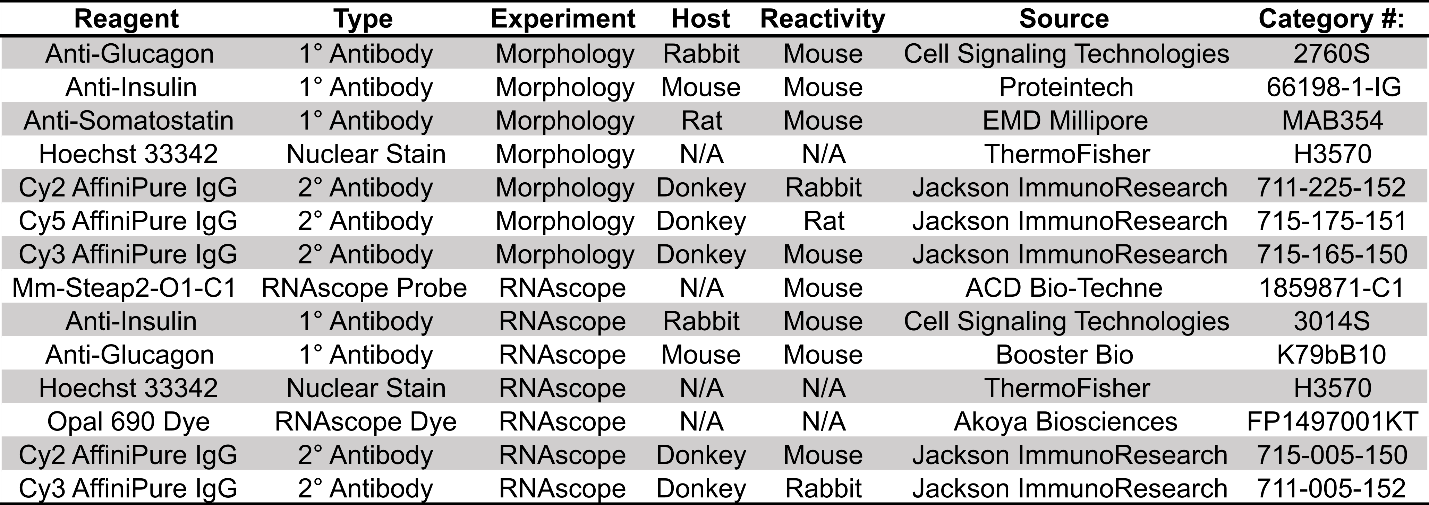

**Supplementary Table 3: List of antibodies and RNAscope probes used.** List of antibodies utilized in immunofluorescence staining of FFPE pancreas sections in morphology and RNAscope experiments. The probes used, type, host, reactivity, manufacturer and category numbers are listed.
